# Cell adhesion promoted by a unique *Shigella* IpaA vinculin- and talin-binding site

**DOI:** 10.1101/329136

**Authors:** Cesar Valencia-Gallardo, Charles Bou-Nader, Daniel Aguilar, Nathalie Carayol, Nicole Quenech’Du, Ludovic Pecqueur, HaJeung Park, Marc Fontecave, Tina Izard, Guy Tran Van Nhieu

## Abstract

During *Shigella* cell invasion, the IpaA effector targets the focal adhesion protein vinculin through three vinculin-binding sites (VBSs). Here, we report that IpaA VBS3 also binds to talin. The 2.5 Å resolution crystal structure indicates that IpaA VBS3 forms a tightly folded α-helical bundle with talin H1-H4, contrasting with bundle unraveling upon vinculin interaction. High-affinity binding of Ipa VBS3 to talin H1-H4 requires a core of hydrophobic residues conserved in vinculin binding and a pair of electrostatic interactions accounting for talin binding specificity. IpaA VBS3 does not bind to talin H1-H5 suggesting the targeting of partially activated stretched talin but not inactive talin. Consistently, IpaA VBS3 labeled filopodial and nascent adhesions and regulated their formation through talin binding. In addition, talin-binding by IpaA VBS3 was required for bacterial capture by filopodia during *Shigella* invasion and for the stabilization of focal adhesions in infected cells. These findings point to the functional diversity of VBSs and a critical role for talin-binding by *Shigella* IpaA VBS3 and possibly talin VBSs in the regulation of adhesion structures.

---

Integrin-mediated cell adhesion critically depends on the anchoring of adhesion receptors to the cytoskeleton, a process regulated by mechanosensing (Riveline et al., 2001). This cytoskeletal anchoring occurs via cytoskeletal linkers, such as talin and vinculin, acting as a molecular clutch coupling adhesion receptors to actin filaments and transducing acto-myosin based contractile forces (del Rio et al., 2009; Fouchard et al., 2014). Following binding of integrin receptors to the extracellular matrix (ECM) and clustering, adhesion structures form and are observed to vary in composition and stability. For ß1-integrins, transient small adhesion structures called focal complexes (FCs) or nascent adhesions are formed at the cell leading edge during the extension of lamellae induced by actin polymerization. FCs mature into focal adhesions (FAs), that are larger, more complex and stable structures. The cytoskeletal linker talin plays a central role in the formation and maturation of focal adhesions (Lagarrigue et al., 2016; Yan et al., 2015). Talin consists of an amino-terminal FERM domain connected to a large rod domain that includes at least two F-actin binding sites (Calderwood et al., 2013; Lagarrigue et al., 2016; Yan et al., 2015). Through its amino-terminal FERM domain, talin binds to the cytoplasmic domain of ß1-integrins and mediates inside-out signaling increasing the affinity of integrins for extracellular matrix proteins (Calderwood et al., 2013; Lagarrigue et al., 2016). Talin-mediated integrin activation requires the Rap1 small GTPase and its effector RIAM, which together with integrins form the MIT (MRL-Integrin-talin) that drives the extension of filopodial “sticky fingers” (Lagarrigue et al., 2016; Lagarrigue et al., 2015). Recruitment of inactive folded talin may directly occur via association between the talin FERM domain and Rap1, or between talin rod domain bundles and two RIAM aminoterminal Talin Binding Sites (TBSs) (Goult et al., 2013b; Zhu et al., 2017). Through these functions, Rap1 and RIAM regulate the initial stages of integrin inside-out signaling, by bringing inactive talin to integrins at the membrane and inducing their activation. Upon activation, talin bridges the cytoplasmic tail of the integrin ß1 subunit with actin filaments (Yan et al., 2015). Stretching of the talin molecule due to actomyosin-dependent contractility leads to the unveiling of VBSs along the talin rod domain, likely responsible for the redistribution of vinculin from a membrane proximal to distal location (Yan et al., 2015). Talin rod VBSs activate vinculin, which, through its tail domain, further tethers actin filaments, leading to increased cell adhesion in response to substrate stiffness (Gingras et al., 2005; Yan et al., 2015). VBSs typically correspond to α-helices of 20-25 residues buried in 4 or 5-helix bundles. In vitro measurements on single molecules indicate that unveiling of VBSs in various talin bundles occurs with a defined hierarchy, which may correspond to different stretched states of talin under increasing forces (Yao et al., 2016). Because of their structure, the talin R2 and R3 bundles are proposed to unfold first in response to stretching forces. Unfolding of talin R2R3 leads to RIAM dissociation and vinculin binding to the exposed VBSs (Goult et al., 2013b). Force measurements on purified bundles indicate that talin bundles such as R1 and R10 unfold at intermediate forces, while other talin bundles such as R7-R9 require higher forces to unfold, within the range exerted at FAs (del Rio et al., 2009; Wang and Ha, 2013; Yao et al., 2014; Yao et al., 2016). The hierarchy of talin VBSs’ exposure in cells, however, may be altered by ligands stabilizing talin bundles or talin folding states. For example, inactive talin is proposed to adopt a dimeric folded conformation, in which the rod domains wrapping around the FERM domains hide the integrin and F-actin binding sites, suggesting an activation step prior to integrin binding (Goult et al., 2013a). While in vitro studies point to the existence of talin conformers containing different exposure states of VBSs, the existence of such conformers in cells remains to be demonstrated.

Bacterial pathogens have evolved remarkable strategies to divert host cell processes. To invade normally non-phagocytic cells, bacteria may express surface ligands to cell receptors, such as integrin or cadherin receptors (Dunn and Valdivia, 2010; Pizarro-Cerda et al., 2012). Alternatively, bacterial invasion may be triggered by the injection of effector proteins into host cells through a type III secretion system (T3SS) (Dunn and Valdivia, 2010; Galan et al., 2014; Valencia-Gallardo et al., 2015). Strikingly, VBSs are also present on type III effectors of the pathogenic bacteria *Chlamydia*, Enteroinvasive *E. coli* and *Shigella* (Thwaites et al., 2015; Valencia-Gallardo et al., 2015). Among these, the *Shigella* IpaA type III effector contains three VBSs located within its 145 carboxy-terminal residues. IpaA VBS1 acts as a super-mimic of talin VBSs that binds to the first helix bundle of the vinculin D1 domain (vD1) and promotes the dissociation of the vinculin head-tail interaction (Izard et al., 2006). VBS2 docks to the second helix bundle of vD1, stabilizing the IpaA VBS1-vinculin interaction and conferring IpaA with the property to bind to vinculin with an unmatched affinity (Tran Van Nhieu and Izard, 2007). Structural characterization of the IpaA VBS3 peptide in complex with vD1 indicates a vinculin activation mechanism similar to that of IpaA VBS1 (Park et al., 2011). Consistently, during *Shigella* invasion IpaA VBS3 cooperates with IpaA VBS1-2 to recruit vinculin at bacterial entry sites (Park et al., 2011). The presence of multiple VBSs in IpaA suggests that multiple tethers to vinculin are required for *Shigella* invasion.

Here, we show that in addition to vinculin, IpaA VBS3 also targets talin by forming a folded globular structure with the talin VBS1 domain containing the H1-H4 helices that differs from the stretched bundle structure expected during full talin activation. By performing structural alignments of talin and IpaA VBSs and mutagenesis, we identified the mechanism conferring to a single alpha-helix VBS the property to bind to talin H1-H4. Our results suggest that IpaA VBS3 targets a semi-activated talin conformer in filopodial and nascent adhesions and that its talin-binding activity is involved in the regulation of filopodial elongation and the formation of FAs.

## Results

### IpaA VBS3 binds to vinculin and talin

We performed a yeast two-hybrid screen using an established human placental cDNA library corresponding to a total of 82.02 million prey clones and a bait containing IpaA1-565 devoid of the VBS1-2 sites (559-633). This screen identified 150 clones representing 16 different genes. Among these, talin was identified with very high confidence in 95 prey clones, with clones corresponding to different open reading frames in talin including the R1 and R10 bundles containing talin VBS1 (482-655) (Papagrigoriou et al., 2004) and VBS3 (1944-1973) (Gingras et al., 2006), respectively (Exp. View Table 1). As expected from the presence of IpaA VBS3 bait, vinculin was also identified as a prey (Exp. View Table 1). This analysis raised the intriguing possibility that IpaA VBS3, in addition to its VBS activity, could also act as a talin-binding site (TBS).

**Table 1.** Non-linear fit values for ITC measurements. N (stoichiometry), K_D_ (affinity constant), δH (cal/mol) enthalpy, -TδS (cal/mol) entropic contribution and ΔG (cal/mol) free enthalpy. IpaA and talin VTBSs binding to talin H1-H4 is exothermic (ΔG<0) and mainly driven by enthalpy (ΔH<0 and ΔH<- TΔS). Talin H1-H4 binding to IpaA peptides VBS_1_ and VBS3 show an important enthalpic and minor entropic contribution. Talin H1-H4 binding to Talin VBS’s shows an important enthalpic contribution and an unfavorable entropic contribution to ΔG. The estimated K_D_ of other talin VBSs for vinculin D1 (K_D_ vD1) was inferred from previous studies (Izard et al., 2006; Izard and Vonrhein, 2004; Park et al., 2011; Yogesha et al., 2012).

| Cell | Titran | $K_D$<br>(Talin H1-H4) | N | $\Delta H$<br>(Kcal/mol) | -TDS<br>(Kcal/mol) | $\Delta G$<br>(Kcal/mol) | $K_D$<br>(vD1) |
| --- | --- | --- | --- | --- | --- | --- | --- |
| Talin H1-H4 | IpaA VBS1 | $15.2 \pm 1.1 \mu M$ | 1.22 | -4.10 | -2.47 | -6.57 | 110 pM |
|  | IpaA VBS2 | - | - | - | - | - | 6.6 nM |
| | IpaA VTBS | $174.5 \pm 18.9 \text{ nM}$ | 0.89 | -6.65 | -2.81 | -9.47 | 54 pM |
| | IpaA VTBS K498A | $163.1 \pm 1.1 \text{ nM}$ | 1.18 | -6.28 | -2.97 | -9.26 | n.d. |
| | IpaA VTBS K498E | $300.3 \pm 2.8 \text{ nM}$ | 0.73 | -12.1 | 3.25 | -8.90 | n.d. |
| | IpaA VTBS R489A K498A | $328.9 \pm 1.9 \text{ nM}$ | 0.69 | -8.68 | -0.16 | -8.84 | n.d. |
| | IpaA VTBS A495K | $2.93 \pm 0.2 \mu M$ | 1.05 | -3.73 | -3.81 | -7.54 | n.d. |
| | Talin VBS46 | $60 \pm 0.8 \text{ nM}$ | 0.92 | -16.97 | 7.20 | -9.77 | 3 nM |
| | Talin VBS6 | $21 \pm 3.4 \mu M$ | 1.26 | -14.34 | 7.97 | -6.37 | n.d. |
| | Talin VBS9 | $1 \pm 0.1 \mu M$ | 1.29 | -14.65 | 6.47 | -8.18 | n.d. |
| | Talin VBS50 | $45 \pm 7.5 \mu M$ | 1.02 | -13.23 | 7.31 | -5.92 | 99 nM |
|  | Tln VBS33 | No binding | - | - | - | - | n.d. |
|  | Tln VBS36 | No binding | - | - | - | - | n.d. |
| A483 (Site 1) | Talin H1-H4 | $115.7 \pm 32.15 \text{ nM}$ | 1 | -6.38 | -3.07 | -9.45 | n.d. |
| A483 (Site 2) | | $4.6 \pm 0.6 \mu M$ | 1 | -6.76 | -5.01 | -7.27 | n.d. |
| A524 (Sites 1 + 2) | | $2.0 \pm 0.5 \mu M$ | 1.52 | -3.97 | -3.78 | -7.76 | n.d. |

To directly show that IpaA VBS3 acted as a Talin Binding site (TBS), we performed native gel shift assays using synthetic peptides (Materials and Methods). As shown in Fig. 1A, IpaA VBS3 formed a complex with talin H1-H4. In contrast, even at high molar ratios, IpaA VBS1 and IpaA VBS2 did not form any detectable complex with talin H1-H4. Consistently, ITC measurements indicated that IpaA VBS3 bound to talin H1-H4 with a high affinity (K_D_ = 174 ± 19 nM), whereas IpaA VBS2 did not show any detectable interaction with talin H1-H4 (Fig. 1B; Table 1). ITC analysis indicated that IpaA VBS1 interacted with talin H1-H4 with much lower affinity than IpaA VBS3 (K_D_ = 15.2 ± 1.1 ;μM; Fig. 1B; Table 1). When native gel shift assays using IpaA 483-633 containing all three VBS1-3 (A483) were performed, a clear migration shift corresponding to a A483-talin H1-H4 complex associated was observed with depletion of free talin H1-H4 (Fig. 1C, arrowhead). In contrast, when IpaA 524-634 containing only IpaA VBS 1-2 (A524) was incubated with talin H1-H4, no migration shift could be detected (Fig. 1C, A524). ITC analysis indicated that talin H1-H4 bound to A483 at one site with an estimated affinity slightly higher than that of IpaA VTBS alone (Exp. View Table 2, K_D_ = 115 ± 32 nM). In addition, a second site with a much lower affinity of 4.6 ± 0.6 μM for A483 binding to H1-H4 could be attributed to IpaA VBS1-2 (Table 1). Consistent with the native gel shift assays, A524 showed a moderate affinity for talin H1-H4 (Exp. View Table 2, K_D_ =2.0 ± 0.5 μM).

**Figure 1.**
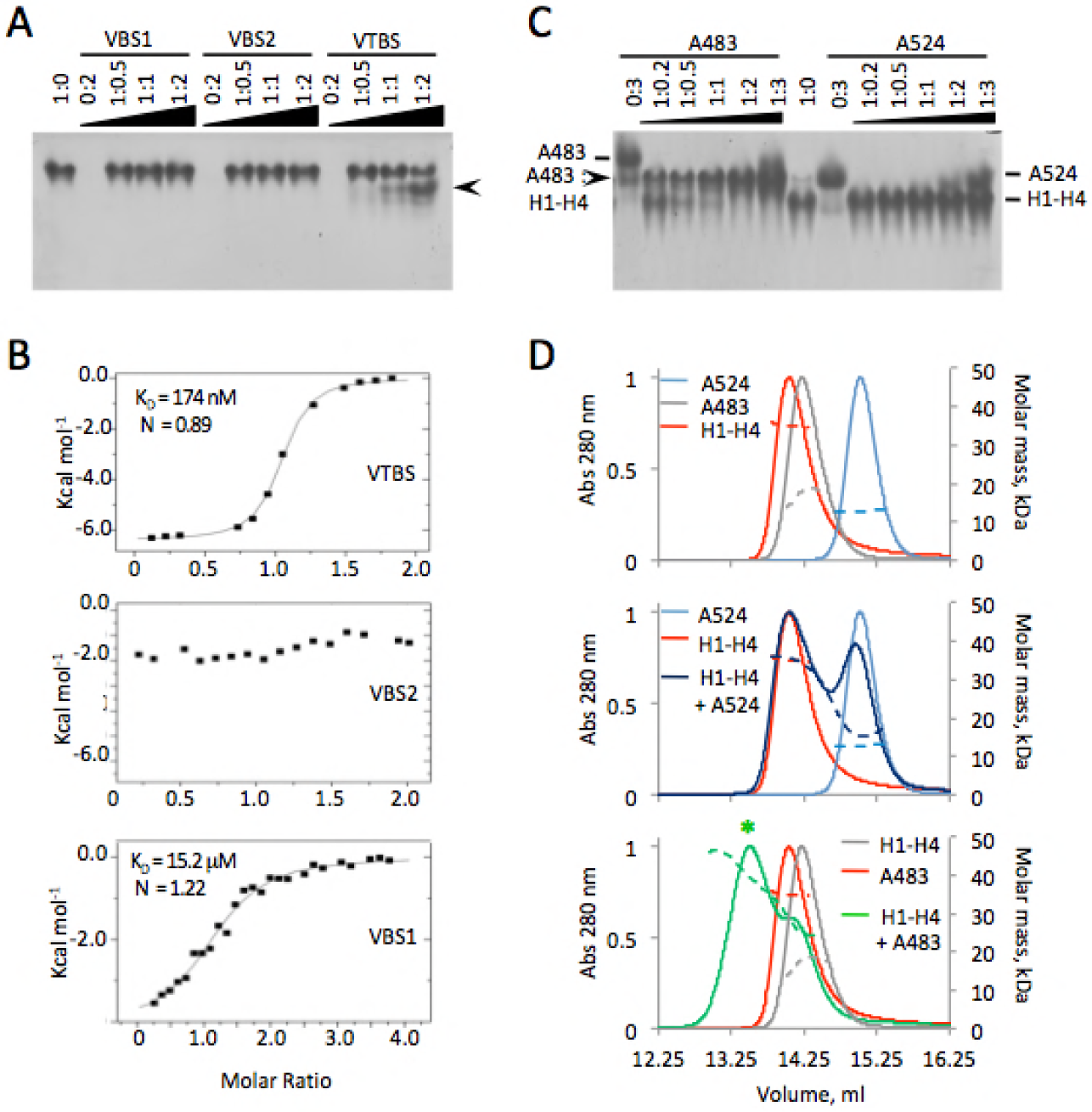
IpaA VBS3 interacts with talin. (A) Analysis of IpaA VBSs and talin H1-H4 interaction using 6-18 % gradient native PAGE. The talin H1-H4: IpaA VBS molar ratio is indicated with 1 corresponding to a final concentration of 25 μM. Arrowheads point to talin H1-H4 and talin H1-H4:IpaA VBS3. (B) Isothermal titration calorimetry (ITC) analysis of the interaction between talin H1-H4 and the indicated IpaA VBSs. The estimated K_D_ are indicated. IpaA VBS2 showed no binding to talin. (C) Native PAGE analysis. The talin H1-H4: IpaA derivative molar ratios are indicated with 1 corresponding to a final concentration of 25 μM. (D) SEC-MALS analysis. Indicated proteins were incubated for 60 min in column buffer prior to SEC analysis using a Superdex 200 10/300 GL increase column. Traces: normalized absorbance at 280 nm of the indicated proteins or complex species in the corresponding color. Dotted lines: molecular mass of the indicated proteins or complexes determined by MALS in the corresponding colors.

**Table 2.** SEC-MALS analysis of the interaction between Talin H1-H4 and IpaA proteins. The molecular mass predicted by the primary sequence (Predicted) of Talin H1-H4 and IpaA proteins and complex was compared to the measured molecular mass (Measured) of these proteins and complex by SEC-MALS at the indicated stoichiometry. The hydrodynamic radius (R_h_) of the complex was determined by dynamic light scattering. Protein concentration was determined by refractometry.

|  | A:H1-H4 | Predicted<br>(kDa) | Measured<br>(kDa) | R<sub>h</sub> (nm) |
| --- | --- | --- | --- | --- |
| A483 | 1 | 17.3 | 17.9 | n.d. |
| A524 | 1 | 12.8 | 12.5 ± 0.7 | n.d. |
| Talin H1-H4 | 2 | 36.4 | 35.2 ± 0.2 | 3.3 ± 0.3 |
| Talin H1-H4 +<br>A483 | 2:1 | 53.7 | 46.8 ± 0.2 | 3.7 ± 0.3 |

Size-exclusion chromatography coupled with multi-angle static light scattering (SEC-MALS) analysis indicated that A483 and A524 were monomers in solution, as indicated by the molecular mass determination (Fig. 1D; Table 2). Of note, H1-H4 behaved as a globular dimer in SEC-MALS analysis (Fig. 1D; Table 2), consistent with the crystal structure of talin rod 482-655 (Papagrigoriou et al., 2004). When mixed prior to analysis, only single species corresponding to either the talin H1-H4 dimer or A524 monomer were recovered, consistent with the native gels and ITC results (Fig. 1D; Table 2). In contrast, a compact globular complex corresponding to one talin H1-H4 dimer bound to a single A483 molecule was observed (Fig. 1D, talin H1-H4 + A483; Table 2). These results are in agreement with the ITC measurements and suggest that A483 strongly interacts with the talin H1-H4 via its VTBS as a first high affinity site. These results indicate that in addition to vinculin, IpaA VBS3, thereafter referred to as IpaA VTBS, also binds to talin with high affinity, a property not shared with other IpaA VBSs.

### IpaA VTBS forms an α-helix bundle with talin H1-H4, mimicking talin H5

We solved the crystal structure of the complex at 2.5 Å resolution (Exp. View Table2). The IpaA VTBS-talin H1-H4 complex was purified using a strategy similar to that used for the IpaA VBS3-vD1 complex (Park et al., 2011). The asymmetric unit contains 6 molecules of talin H1-H4 organized into dimers as observed in SEC-MALS (Expanded View Fig. 1A). Each talin H1-H4 molecule shows a clear additional density corresponding to the IpaA VTBS peptide in vicinity of mainly the H2 and H4 helices (Expanded View Fig. 1B). All talin H1-H4 chains in the asymmetric unit were very similar with pairwise RMSDs in the range 0.28-0.64 Å over 140 Cα. The interface area between IpaA VTBS and talin H1-H4 is 849 ± 26 Å^2^ and involve mainly hydrophobic residues located in talin α-helices H2, H3 and H4 (Figs. 2A and Expanded View Fig. 2C). In addition, electrostatic interactions, either direct, between K498 and E621, or indirect with R489 pointing within a negatively charged region further strengthen IpaA VTBS binding and positioning in the talin H2-H4 groove (Figs. 2A, 2B).

**Figure 2.**
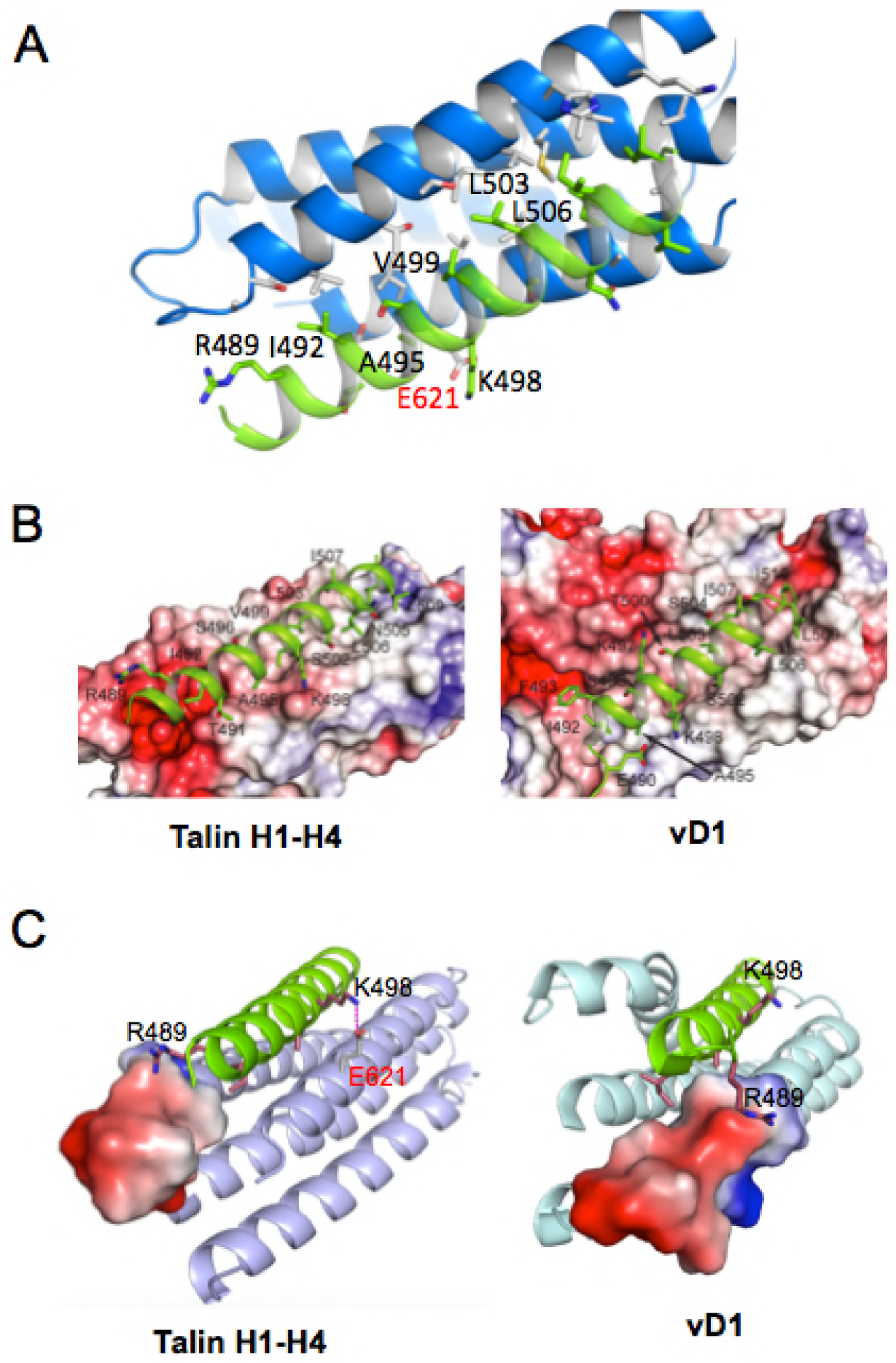
IpaA VTBS forms a compact fold with talin H1-H4. (A) X-ray structure of the talin H1-H4:IpaA VTBS complex. Blue: talin polypeptide chain. White: interacting residues. Green: bound IpaA VTBS with interacting residues in green. VTBS residues are annotated according to full length IpaA. (B, C) Molecular detail of the interaction between IpaA VTBS with talin H1-H4 (B) and vD1 (C). IpaA VTBS is shown in cartoon with its relevant interacting residues. Talin H1-H4 is illustrated by its vacuum electrostatic potential generated with APBS [38]. Molecular detail of the interaction between IpaA VTBS with vinculin D1 domain. IpaA VTBS is shown in cartoon with its relevant interacting residues. Vinculin D1 is illustrated by its vacuum electrostatic potential generated with APBS [38].

Comparison of talin H1-H4 in its IpaA VTBS bound state onto talin H1-H5 (Expanded View Fig. 2) revealed a remarkable superimposition of VTBS with H5. Indeed, I492, A495, V499, L506 and I507 of IpaA VTBS involved in hydrophobic interaction with talin H1-H4 are homologous to L637, A640, V644, L651 and L652 in the talin α-helix H5, nesting in the identical hydrophobic groove (Expanded View Figs. 2A and 2B). Furthermore, as shown in Expanded View Fig. 2C, a strong similarity in terms of polarity is observed between talin H5 and IpaA VTBS, with R489 in IpaA VTBS corresponding to R634 in talin H5 engaged in the same polar interaction with the talin H2-H3 loop.

Structural comparison of IpaA VTBS in its talin-bound and vinculin-bound state (PDB 3rf3, Figs. 2B, 2C) indicated that the same IpaA VTBS hydrophobic residues were involved in binding to talin H1-H4 and vD1. The IpaA VTBS-vinculin interaction surface area, however, is wider (1150 Å^2^) than that of talin H1-H4 and comprises IpaA VTBS residues F493, K497 and E490, which are not involved in talin H1-H4 recognition (Figs. 2B, 2C). Strikingly, IpaA K498 was found to be specific for interaction with talin since it targets talin E621 and does not establish interaction with vD1 (Figs. 2B, 2C).

These results indicate that IpaA VTBS interacts with the talin H1-H4 bundle in a mode that does not imply the major unfolding and exposure of the talin H4 helix required for binding to the canonical vD1 talin-binding domain. Instead, binding of IpaA VTBS to talin H1-H4 mimics that of talin H5 and targets a talin conformer containing a compact H1-H4 folded bundle.

### Contact residues in VTBSs determining binding specificity for talin

Synthetic peptides were generated to confirm the role of IpaA polar and hydrophobic residues in talin H1-H4 binding. Specifically, we tested the effects of charge removal at R489 and K498 by alanine substitution, charge inversion by substitution of K498 to glutamic acid, and substitution of hydrophobic A495 by a positively charged lysine residue. The latter A495K mutation was designed from our structural alignment indicating that such substitution could explain the lack of talin binding for IpaA VBS2 (Fig. 3A; Table 1). As indicated by ITC analysis, the K498A substitution did not lead to a significant decrease in talin binding affinity of IpaA VTBS, possibly because of the complementary charge contribution by adjacent K497 (Fig. 3A; Table 1). The single charge inversion K498E, however, led to a defect in talin H1-H4 binding with a 1.7-fold decrease in the determined K_D_ (K_D_ = 300 ± 2.8 nM). A similar decrease was observed for the charge R489A K498A suppressing double mutations with a K_D_ = 328 ± 1.9 nM. Consistent with a critical role for the hydrophobic interactions, the A495K substitution led to a drastic defect in IpaA VTBS binding to talin, with a 17-fold decrease in K_D_ (K_D_ = 2.93 ± 0.3 μM). These results were further confirmed in native gel shift experiments showing that, as opposed to IpaA VTBS, IpaA VTBS 498E and IpaA VTBS A495K did not induce any detectable shift in the presence of talin H1-H4 (Fig. 3B). In contrast, shifts were still observed in the presence of vD1, indicating that the mutations did not prevent complex formation with vinculin, although higher peptide concentrations were required for the A495K substitution consistent with a partial effect of this substitution on vinculin binding (Fig. 3B).

**Figure 3.**
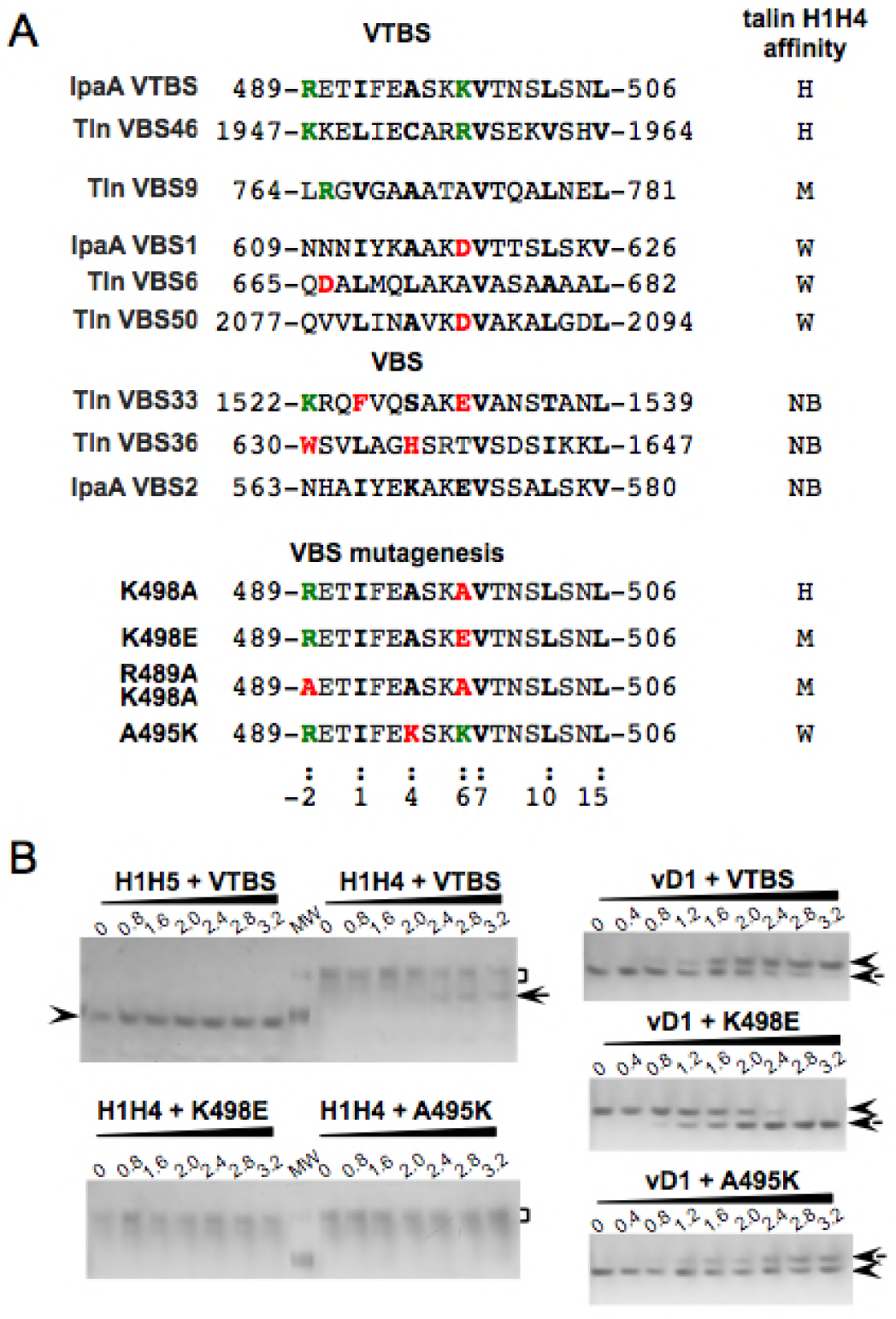
Structural alignments of talin and IpaA VBSs. (A) Sequence alignment of IpaA VTBS and talin VBSs. The K_D_ for talin H1-H4 was determined by ITC analysis and used to classify the VTBS in high- (H), moderate- (M), or weak-affinity (W) and Non-binders (NB). The contact residues equivalent to R489 and K498 of IpaA VTBS are indicated. The residues involved in hydrophobic interactions are shown in bold. Modification of these residues was annotated as follows: green: conserved; red: charge change / large change. (B) Analysis of IpaA VTBS, VTBS A495K and VTBS K498E variants’ interaction with talin H1-H4 (B) or vD1 (C) using 6-18 % gradient native PAGE. The talin H1-H4:IpaA VTBS or VD1: IpaA VTBS molar ratio is indicated with 1 corresponding to a final concentration of 25 μM. Arrowheads point to talin H1-H4 and talin H1-H4:IpaA VTBS3. The arrowhead and bracket point to uncomplexed talin H1-H5, talin H1-H4, or vD1. The arrows point to talin H1-H4: IpaA VTBS, vD1: IpaA VTBS, or vD1: IpaA VTBS A495K complexes. (C)The dotted arrow points to vD1: IpaA K498E complex. MW: molecular weight standards.

To investigate whether endogenous single helix talin VBSs could use a similar mechanism as IpaA VTBS to bind to talin, we measured the affinity of synthetic peptides corresponding to talin VBS6, VBS9, VBS33, VBS36, VBS46 and VBS50 for talin H1-H4 (Table 1). Talin VBSs could be divided in four classes according to their affinity for talin H1-H4 determined by ITC: while VBS33 and VBS36 did not show detectable binding, VBS6, VBS50 and VBS9 showed weak or moderate affinity for talin H1-H4 (Table 1). Talin VBS46, however, interacted with talin H1-H4 with an affinity comparable to that of IpaA VTBS, with an estimated K_D_ = 60 ± 0.8 nM (Table 1). No obvious correlation was observed between the respective affinity of these VTBSs for talin H1-H4 and their reported affinity for vinculin D1 domain (Table 1, vD1). While consistently displaying high affinity for vD1, VTBSs bound to talin H1-H4 with a broad range of K_D_ values (Table 1).

To identify residues conferring the property of these VBS single alpha-helices to bind to talin, we aligned the amino acid sequences of IpaA and talin VBSs (Fig. 3A). This analysis indicated that contact residues in IpaA VTBS involved in hydrophobic interactions with vinculin or talin are generally conserved in talin VBS46, conferring the minimal hybrid vinculin-talin binding property (Fig. 2B). Interestingly, talin VBS46 (talin residues 1947-1964) and IpaA VTBS which both bind to talin H1-H4 with high affinity (K_D_ of 60 and 175 nM, respectively) share a pair of positively charged residues at positions “-2” (IpaA Arg 489 and talin Lys 1947) and “7” (IpaA Lys 498 and talin Arg 1956). These positively charged residues are identified in our structure to be a key in positioning and stabilizing IpaA VTBS in the talin H2-H4 groove through electrostatic interactions and are likely essential to confer high affinity binding (Fig. 2C). Consistently, talin VBS9 (talin residues 764-781) which binds to talin H1-H4 with a K_D_ in the low micromolar range, retains an Arg residue at position “-2”, but has an alanine at position “7” that might lead to reduced affinity (Table 1, Fig. 3A). Furthermore, two weakly binding α-helices, namely IpaA VBS1 (IpaA residues 609-626) and talin VBS50 (talin residues 2077-2094) that have lower binding affinities with K_D_ in the 15-45 μM range, show a loss of charge at position “-2” (Asn 609 for IpaA and Gln 2077 for VBS50) combined with a charge inversion at position “7” (Asp 618 for IpaA and Asp 2086 for VBS50) that might further reduce affinity via electrostatic clash. Talin VBS33, VBS36 and IpaA VBS2 that do not bind talin H1-H4 present a combination of charge inversion and substitutions in the hydrophobic interface, suggesting that the latter also plays a role in specifying talin binding (Table 1, Fig. 3A). Scrutinizing of changes in this hydrophobic core indicates that while hydrophobic residues at positions 8, 12 and 15 are conserved for all these VTBSs and VBSs, at least one substitution of hydrophobic to polar residues is observed at positions 1 and 4 for VBSs (Fig. 3A).

Together, our structural data, comparison of the IpaA and talin VBS sequences, as well as ITC analysis are fully consistent and point to specific hydrophobic interactions at positions 1 and 4 and electrostatic interactions at positions −2 and 7 as being important for talin H1-H4 binding. This is illustrated by the structures of IpaA VTBS in complex with vD1 or H1-H4 in Figs. 2B, C. In the vinculin-IpaA structure, IpaA residue K498 at position 7 is solvent exposed, suggesting that a positively charged residue is important for binding to talin but not to vinculin (Fig. 2C, Exp. View Fig. 2). Of note, the aminoterminal residues Thr 489 to Thr 487 of IpaA VTBS form an additional helical turn when bound to talin H1-H4 as opposed to in the vD1 complex. This suggests that IpaA VTBS folds differently upon binding to talin or vinculin, further tuning recognition specificity.

### IpaA VTBS labels talin- and vinculin-containing adhesions structures

TBSs corresponding to single α-helices in RIAM and VLC1/KANK have been reported to bind to the folded talin bundles R2R3 and R8, respectively, consistent with targeting of inactive talin (Goult et al., 2013b; Zacharchenko et al., 2016). In contrast, our evidence suggests that IpaA VTBS targets an active conformer of talin because of its binding to talin H1-H4 but not talin H1-H5. Remarkably, IpaA VTBS forms a folded helical bundle that differs from the “opened” configuration reported to interact with vinculin (del Rio et al., 2009; Papagrigoriou et al., 2004). These results suggest that IpaA VTBS targets a semi-stretched conformer of talin that is different from inactive folded talin and a fully stretched conformer.

To test the dual talin-vinculin binding activity of IpaA VTBS and targeting of a specific talin conformer, we took advantage of its high affinities for vinculin and talin to localize its target in cell-labeling experiments. We generated a GFP fusion to IpaA VTBS and analyzed its localization following cell transfection. As shown in Fig. 4A, GFP-IpaA VTBS labeled FAs of cells plated onto fibronectin-coated glass coverslips, showing a strict co-localization with talin and vinculin. As expected, the number of GFP-IpaA VTBS labeled-adhesion structures decreased in siRNA-mediated vinculin-, talin- or talin and vinculin-depleted cells (Figs. 4B-D). In talin-depleted cells, GFP-IpaA VTBS co-localized with vinculin-labeled structures consistent with its vinculin binding activity (Fig. 4B, siRNA vinculin). Conversely, in vinculin-depleted cells, GFP-IpaA VTBS co-localized with talin structures consistent with talin binding (Fig. 4B, siRNA vinculin). These results are consistent with IpaA VTBS binding to talin and vinculin.

**Figure 4.**
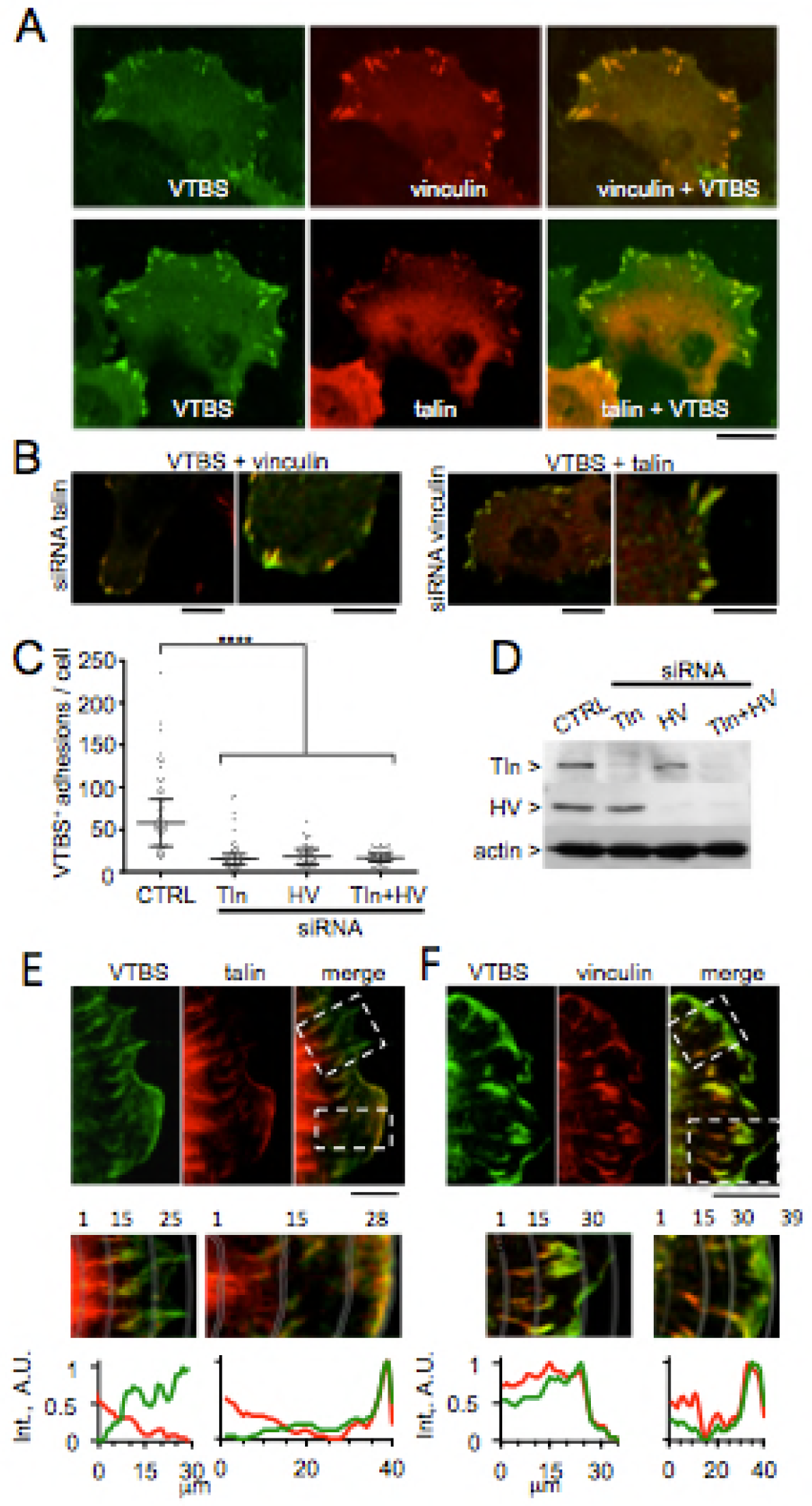
IpaA VTBS labels focal adhesions. (A, B) Representative confocal micrographs of HeLa cells transfected with the indicated fluorescent markers, fixed and processed for fluorescence staining. Green: GFP-IpaA VTBS; red: vinculin or talin; cyan: actin. (B) Cells treated with the indicated siRNA prior to transfection with IpaA VTBS. Note the co-localization of IpaA VTBS with vinculin and talin in cells treated with anti-talin and anti-vinculin siRNAs, respectively. Scale bar = 5 µm. (C) The number of FAs of control and siRNA treated cells was quantified (Methods) and compared using a Kruskal-Wallis test with Dunn’s multiple comparisons test (****: p < 0.001). (D) HeLa cells were treated with anti-talin and/or anti-vinculin siRNA for 16 hours at a concentration of 40nM. Crude lysates were analyzed by anti-talin, anti-vinculin and anti-actin Western blotting. Representative Western-blot analysis using antibodies against the indicated proteins. In typical experiments, vinculin and talin levels were decreased by at least 80 % in cells treated with the corresponding siRNA. (E, F) MEF cells plated on 15-kPa stiffness substrates were transfected, fixed and processed for fluorescence staining. Samples were analyzed by confocal microscopy. Cells transfected with: GFP-IpaA VTBS (VTBS), talin-mCherry, and vinculin-mCherry. Representative micrographs are shown. Scale bar: 5 μm. Lower panels correspond to higher magnifications of the insets boxed in top panels. Bottom: scans of the integrated fluorescence intensity of concentric area in dotted lines in the corresponding fluorescence micrographs in adhesion structures and lamellae shown in the top panels. Note that IpaA VTBS co-localizes with talin and vinculin at lamellae‘s edge and cortical adhesions, but not with talin-containing fibrillar structures or vinculin-containing larger adhesions in the interior part of the cell.

To favor the formation of adhesion structures subjected to low forces relevant for a partial talin activation state, we plated cells on 15 kPa stiffness substrate coated with fibronectin (Materials and Methods). As shown in Figs. 4E, F, IpaA VTBS labeled vinculin and talin-containing adhesions at the cell periphery, including at the base of filopodia as well as the edge of lamellae. Adhesion structures located towards the cell interior, however, were less enriched in IpaA VTBS, consistent with IpaA VTBS labeling nascent adhesions (Figs. 4E, F) (Beningo et al., 2001; van Hoorn et al., 2014).

### Talin-binding by IpaA VTBS is required for targeting of adhesions

To induce the formation of adhesions, cells were lifted up by trypsinization and analyzed following short time periods after replating (Material and Methods). These replating experiments confirmed the labeling of nascent adhesions by GFP-IpaA VTBS that was observed when cells were plated on low stiffness substrate, with the labeling of talin and vinculin-containing peripheral adhesions as well as prominent adhesions 20 min following replating (Fig. 5, VTBS). To test the role of talin binding by GFP-IpaA VTBS in adhesion targeting, cells were transfected with the mutated IpaA VTBSs containing the A495K or K498E substitutions. As shown in Figs. 5A-C, these mutated IpaA VTBSs impaired for talin binding failed to localize to adhesion structures, with a 6- and 7.8-fold reduction in the number of GFP-IpaA VTBS-positive adhesions per cell for IpaA VTBS A495K and K498E, respectively, compared to parental IpaA-VTBS (Figs. 5A-C). In addition, the size of talin-containing adhesions was significantly reduced in GFP-IpaA VTBS A495K and K498E transfected cells (Fig. 5D). Similarly, GFP-IpaA VTBS A495K and K498E affected the size of vinculin-containing adhesion structures (Fig. 5E).

**Figure 5.**
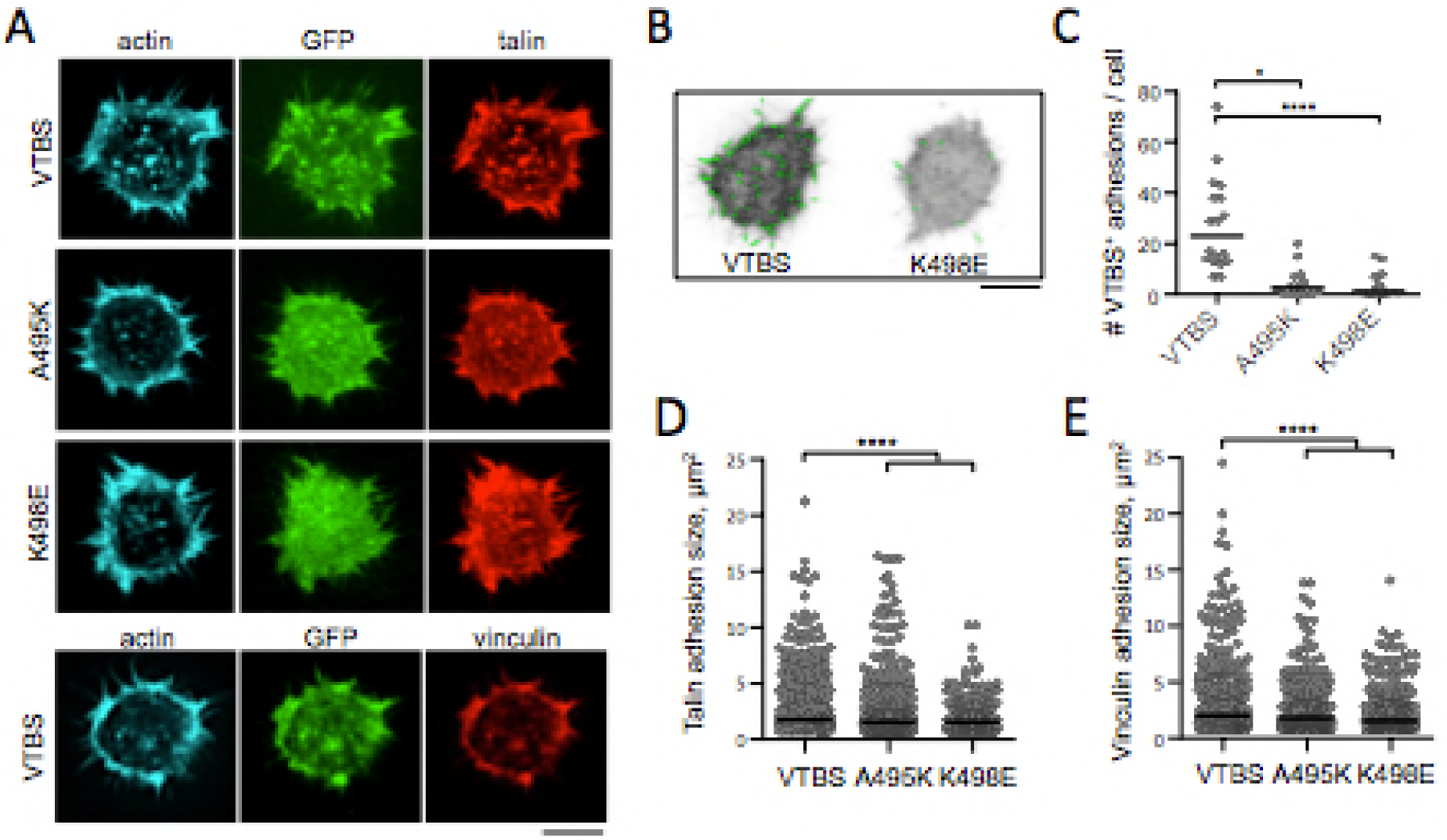
Talin-binding by IpaA VTBS promotes focal adhesions formation. Cells were trypsinized and replated onto fibronectin-coated glass coverslip for 15 min prior to fixation and processing for fluorescence staining. Samples were analyzed by confocal microscopy. Cells transfected with: GFP: GFP-IpaA VTBS (VTBS), A495K, or K498E variant and talin-mCherry (talin) or mCherry-vinculin (vinculin). (A) Representative fluorescence micrographs of cell basal planes. Scale bar: 5 μm. (B, D) Wavelet Spot Detection (green) of adhesion structures in replated cells (Materials and Methods). (C) Distribution of cells as a function of the number of detected VTBS-labeled adhesions. Median ± MAD: VTBS: 23 ± 10.5 (18 cells, N=3); A495K: 2.5 ± 2.5 (14 cells, N=3); K498E: 1 ± 1 (17 cells, N=3) Dunn‘s test. *: p < 0.05. ****: p < 0.001. (E) Size distribution of FAs for talin-mCherry transfected cells. Median ± MAD: VTBS: 1.79 ± 0.76 (774 FAs, N=3); A495K: 1.53 ± 0.51 (568 FAs, N=3); K498E: 1.53 ± 0.51 (383 FAs, N=3). Dunn‘s test. *: p < 0.05. ****: p < 0.001. (F) Size distribution of FAs for mCherry-vinculin transfected cells. Median ± MAD: VTBS: 2.04 ± 0.76 (629 FAs, N=3); A495K: 1.79 ± 0.76 (595 FAs, N=3); K498E: 1.53 ± 0.51 (480 FAs, N=3). Dunn‘s test. *: p < 0.05. ****: p < 0.001.

Together, these results indicate that IpaA VTBS targets nascent adhesions. The effects of IpaA VTBS A495K and K498E mutants suggest that talin binding by IpaA VTBS regulates adhesion formation.

### IpaA VTBS labels filopodial distal adhesions enriched in talin

In addition to the labeling of cell adhesions by IpaA VTBS, we observed labeling of filopodial adhesions. First, we found that in replating experiments, talin was enriched in adhesions in the filopodial shaft distal moiety while vinculin preferentially localized at the filopodial shaft proximal moiety and base (Fig. 6A). To quantify this, we used a semi-automated procedure based on wavelet-spot detection to identify filopodial adhesions and to determine the location of filopodial vinculin and talin adhesions relative to the cell body (Materials and methods). As shown in Figs. 6A, B, talin adhesions were more distal to the cell body compared to vinculin clusters with a median distance ± MAD of 2.5 ± 1.8 μm and 1.7 ± 1.3 μm, respectively (Fig. 6B). We next analyzed the location of VTBS-labeled adhesions respective to that of talin- and vinculin-labeled adhesions in filopodia. Due to the weak signal of filopodial adhesions and lack of compatible reagents, we did not succeed in performing vinculin, talin and VTBS triple-labeling. We thus performed dual labeling of filopodial adhesions and determined their distance relative to the cell body (Materials and Methods; Fig. 6C). As shown in Figs. 6A and 6C, GFP-IpaA VTBS was found associated with filopodial distal talin adhesions, whereas association with vinculin adhesions was observed in more proximal location in filopodia, with a median distance for VTBS + talin adhesions of 2.8 ± 1.5 μm compared to 1.4 ± 0.9 μm for VTBS + vinculin adhesions (Fig. 6C). Moreover, VTBS + talin adhesions were located more distally than talin + vinculin adhesions in filopodia, suggesting the formation of distinct complexes (Fig. 6C). Our structural studies predict that IpaA VTBS and vinculin recognize different talin conformers that could account for distinct vinculin-talin and IpaA-talin complexes. Consistent with this, vD1 used as a canonical talin-binding domain labeled filopodial clusters that were more proximal to the cell body than those labeled with VTBS, with a median distance to the cell body of 2.5 ± 1.8 μm and 3.1 ± 2.2 μm, respectively (Fig. 6D). Of note, direct comparison in cells co-expressing mCherrry-vD1 and GFP-IpaA VTBS was not possible because of cytosolic sequestration likely due to interaction between the two constructs.

**Figure 6.**
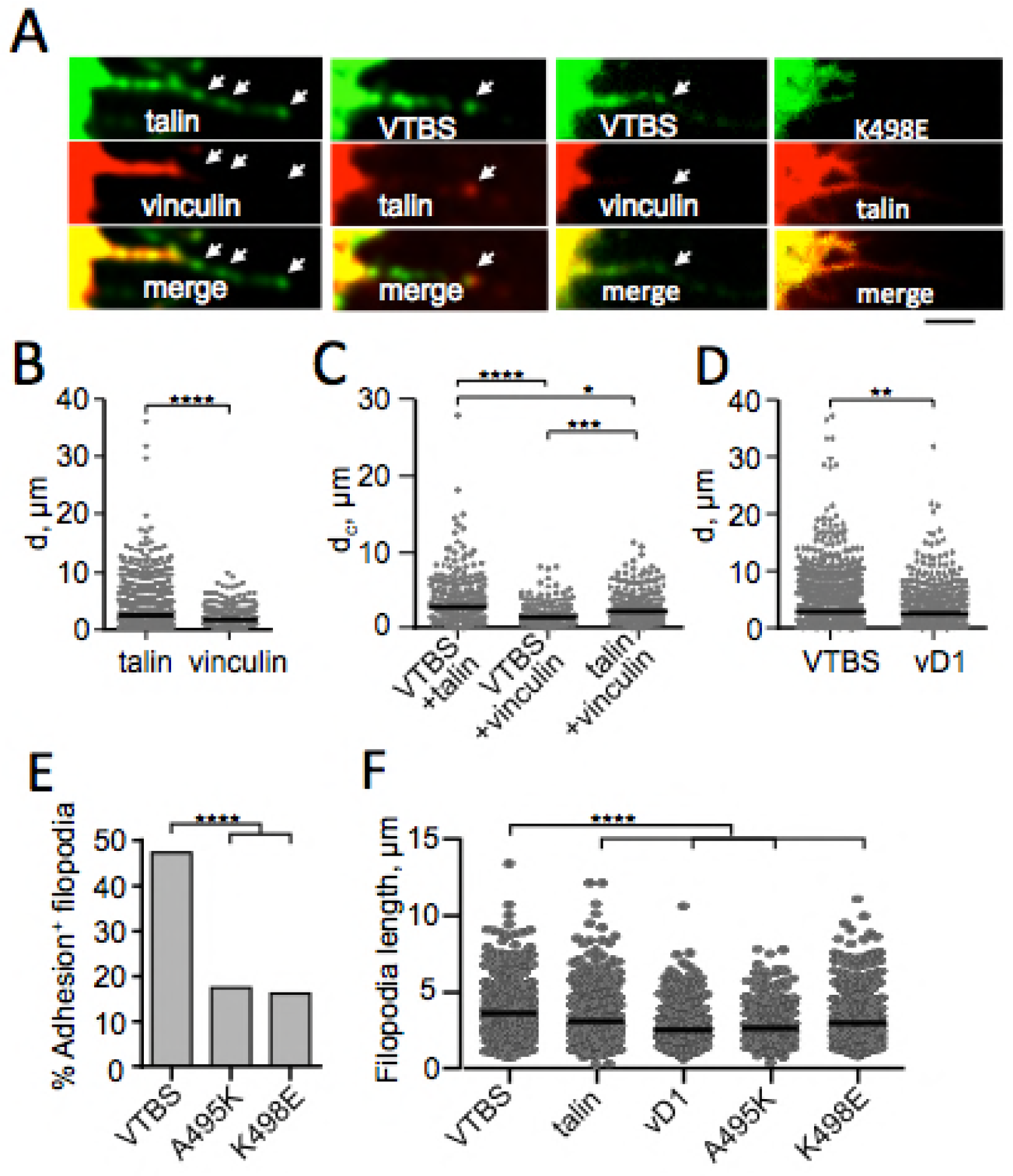
IpaA VTBS targets talin-containing filopodial adhesions. Cells transfected with vinculin-mCherry and: talin-mCherry, GFP-IpaA VTBS ore K498E variant. (A) Representative confocal micrographs. Scale bar = 5 μm. (B, D) Distribution of the distance relative to the cell body (d) of filopodial clusters labeled with the indicated marker in cells co transfected with: GFP-talin and vinculin-mCherry (B); VTBS or vD1 and vincuiin-mCherry (D). Median ± MAD: talin: 2.48 ± 1.81 (858 clusters, N = 3); vinculin: 1.71 ± 1.31 (252 clusters, N = 3); VTBS: 3.08 ± 2.22 (1008 clusters, N = 3); vD1: 2.53 ± 1.79 (537 clusters, N = 3). Mann-Whitney test. **: p < 0.01. ****: p < 0.001. (C) Distribution of the distance relative to the cell body (d) of filopodial clusters co-labeled with the indicated markers in co-transfected cells. Median ± MAD: VTBS+talin: 2.86 ± 1.49 (285 clusters, N =3); VTBS+vinculin: 1.43 ± 0.88 (128 clusters, N = 3); talin+vinculin: 2.32 ± 1.36 (209 clusters, N = 3). Dunn‘s test. *: p < 0.05. ***: p < 0.005. ****: p < 0.001. (E) Percent of filopodia showing VTBS clusters. VTBS (563 filopodia, N = 3); A495K (375 filopodia, N = 3); K498E (332 filopodia, N = 3). Pearson’s χ^2^ test. **: p < 0.001 (F) Dot plot of filopodial lengths. Bar: median MAD. VTBS: 34.5 3.6 μm (656 filopodia, N = 4); talin: 24.1 4.4 μm (435 filopodia, N = 4); vD1: 21.0 3.6 μm (336 filopodia, N = 3); A495K: 2.6 0.9 μm (345 filopodia, N = 3); K498E: 2.9 1 μm (434 filopodia, N = 3); Dunn‘s test. **: p < 0.01. ***: p < 0.005. ****: p < 0.001.

Together, these results suggest that IpaA VTBS labels a talin conformer in filopodial distal adhesions distinct than the one recognized by the vD1 canonical talin-binding domain.

### IpaA VTBS regulates filopodial extension through its talin-binding activity

To test the role of talin-binding by IpaA VTBS in the targeting of filopodial distal adhesions, we analyzed the localization of the GFP-IpaA VTBS A495K and K498E mutants. As shown in Figs. 6A, E, as opposed to GFP-IpaA VTBS, the mutated VTBS variants impaired for talin binding showed a decreased ability to label filopodial adhesions with a 2.7-fold decrease in filopodia containing large adhesions (Fig. 6E). We next investigated the functional consequence of IpaA VTBS binding to filopodial adhesions.

In previous works, binding of vD1 to activated talin was shown to inhibit refolding and stabilize the stretched talin conformation and focal adhesions (Atherton et al., 2015; Yao et al., 2014), we therefore expected binding of IpaA VTBS to favor filopodial extension by stabilizing filopodial adhesions. Consistently, quantification indicated that filopodia in GFP-IpaA VTBS transfectants were longer than those observed in cells transfected with GFP-talin with a median length ± MAD of 4.1 ± 1.9 μm and 3.4 ± 1.9 μm, respectively (Fig. 6F). Remarkably, cells transfected for the GFP-IpaA VTBS A495K and K498E showed filopodia that were significantly shorter than cells transfected with GFP-IpaA VTBS with a median length of 2.6 ±0.8 μm and 2.9 ± 1 μm respectively, and comparable to the filopodial length determined in cells transfected with vD1 that binds to fully stretched talin (Fig. 6G, median 2.5 ± 1.8 μm).

Together, these results suggest that through talin binding, IpaA VTBS stabilizes filopodial adhesions thereby favoring their extension. The role of IpaA VTBS in the regulation of both filopodial and nascent adhesions further supports the link reported between these adhesion structures (Hoffmann and Schafer, 2010; Jacquemet et al., 2015; Partridge and Marcantonio, 2006).

### A role for talin-binding by IpaA VTBS in bacterial capture by filopodia

*Shigella* invasion of epithelial cells is preceded by its capture by filopodia (Romero et al., 2011). To determine if filopodial adhesions targeted by IpaA VTBS talin are involved in bacterial capture, we challenged cells transfected with GFP-IpaA VTBS with *Shigella* and challenged cells for 15 min, a time point corresponding to the early steps of invasion where filopodial capture occurs (Romero et al., 2011). As shown in Fig. 4A, GFP-IpaA VTBS and talin structures could be observed at the tip of filopodia capturing bacteria (Fig. 7A, arrowhead). Consistent with a role during filopodial capture and retraction, this IpaA-VTBS-labeled talin structure was detected at the tip of 84.6 % of filopodia contacting bacteria (13 filopodia, N = 3), but only in 12.8 % of filopodia devoid of bacteria (304 filopodia, N = 3). To test the relevance of IpaA and, talin-binding by IpaA VTBS during filopodial capture, cells were challenged with *ipaA* mutant strains complemented with various IpaA constructs. As shown in Fig. 7B, deletion of all IpaA VBSs resulted in a five-fold reduction in the number of bacterial capture compared to full-length IpaA. While deletion of IpaA VBS1-2 did not affect filopodial capture, deletion of IpaA VTBS reduced by 2.4-fold filopodial capture (Fig. 7B). Introduction of the A495K or K498E mutations that decrease talin binding of IpaA VTBS in IpaA ΔVBS1-2 reduced filopodial capture to an extent similar to that observed for IpaA ΔVTBS (Fig. 7B). Together, these results are consistent with a role for IpaA VTBS in the regulation of filopodial adhesion, and indicate a role for talin binding by IpaA VTBS during filopodial capture.

**Figure 7.**
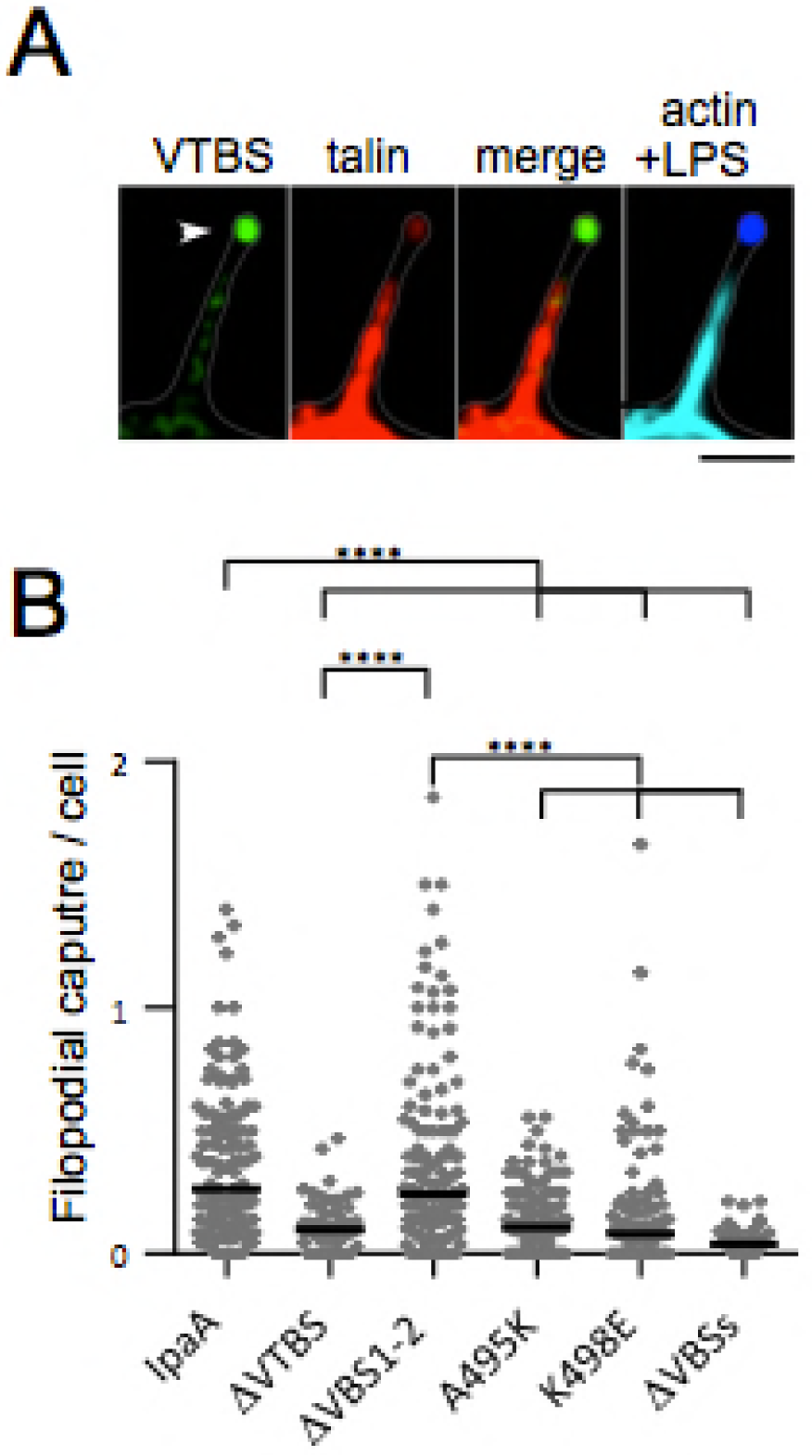
A role for Talin-binding by IpaA VTBS in bacterial capture by filopodia. HeLa cells were challenged with *Shigella* strains for 15 min to allow for filopodial capture. When indicated, cells were transfected prior to bacterial challenge. Samples were fixed and processed for immunofluorescent staining. WT: wild-type *Shigella*. *ipaA*: *ipaA* mutant. *ipaA* mutant complemented with: A: full-length IpaA; ΔVTBS: IpaAΔVTBS; ΔVBS1-2: IpaA ΔVBS1-2; A495K: IpaAΔVBS1-2 A495K; K498E: IpaAΔVBS1-2 K498E; ΔVBSs: IpaA ΔVBS1-3. (A) Representative micrographs. Cells co-transfected with IpaA-VTBS and talin-mCherry and challenged prior to bacterial challenge. Green IpaA VTBS; red: talin; blue: bacteria; cyan: F-actin. Scale bar = 5 μm. Talin and GFP-IpaA VTBS are detected at the filopodial tip where bacterial capture occurs, as well as in bacterial coats in invasion foci induced by WT *Shigella* but not the *ipaA* mutant. (B) The number of bacteria associated with filopodia per cell was scored. A (1739 cells, N = 5); ΔVTBS (1258 cells, N = 2); ΔVBS1-2 (1728 cells, N = 5); A495K (1316 cells, N = 4); K498E (1291 cells, N = 4); ΔVBSs (909 cells, N = 2). Mann-Whitney test. ****: p < 0.001.

### Talin is required for *Shigella* invasion and is recruited in an IpaA-dependent manner at bacterial invasion sites

We next wanted to determine the role of talin binding by IpaA VTBS in the formation of bacterial-induced adhesion structure during *Shigella* invasion

When analyzed by immunofluorescence microscopy, talin was detected in coat-structures around internalized bacteria as early as 10 min-incubation at 37°C (Expanded View Figs. 3A, B, arrowhead), concomitantly with the depolymerization of actin in membrane ruffles and its coalescing around internalized bacteria (Figs. 7A, B and Expanded View Figs. 3A, B). GFP-IpaA VTBS was highly enriched in talin-containing coat-structures surrounding WT *Shigella* but not the *ipaA* mutant (Expanded View Figs. 3C, D).

Cell treatment with anti-talin siRNA did not impair actin polymerization at invasion sites induced by wild-type *Shigella* but actin coat-structures surrounding invading bacteria seldom formed, reminiscent of foci induced by an *ipaA* mutant (Figs. 7A, B). As a result, the percent of internalized bacteria was reduced by 4-fold in anti-talin siRNA treated cells compared to control cells (Fig. 8C). Consistent with a role for IpaA in recruiting talin, actin polymerization foci induced by an *ipaA* mutant strain failed to form talin-coats (Figs. 8D-E, Expanded View Figs. 3E-F). When cells were challenged with *ipaA* mutant strains complemented with IpaA derivatives, talin coat-structures were not detected for the *ipaA* mutant complemented with with vector alone or IpaA deleted for all its VBSs (Figs. 8D, E). Complementation of the *ipaA* mutant with full length IpaA, IpaA ΔVBS1-2 or IpaA ΔVTBS, restored the formation of foci with talin coat structures (Fig. 8E). However, a significant decrease in the percentage of foci forming talin coat structures was observed for the *ipaA* mutant strains expressing IpaA ΔVBS1-2 or IpaA ΔVTBS compared to full length IpaA, with 19 ± 0.7 % and 25 ± 5 % relative to 52 ± 6.7 %, respectively (Fig. 8E). These results indicate that talin recruitment could occur via IpaA VTBS, or via IpaA VBS1-2 possibly through vinculin-talin interactions. Consistently, *ipaA* mutant strains complemented with IpaA ΔVBS1-2 containing the A495K or K498E mutations that have reduced binding for talin but still bind to vinculin, recruited talin coats at a similar extent than the IpaA VBS1-2 complemented strain (Fig. 8F).

**Figure 8.**
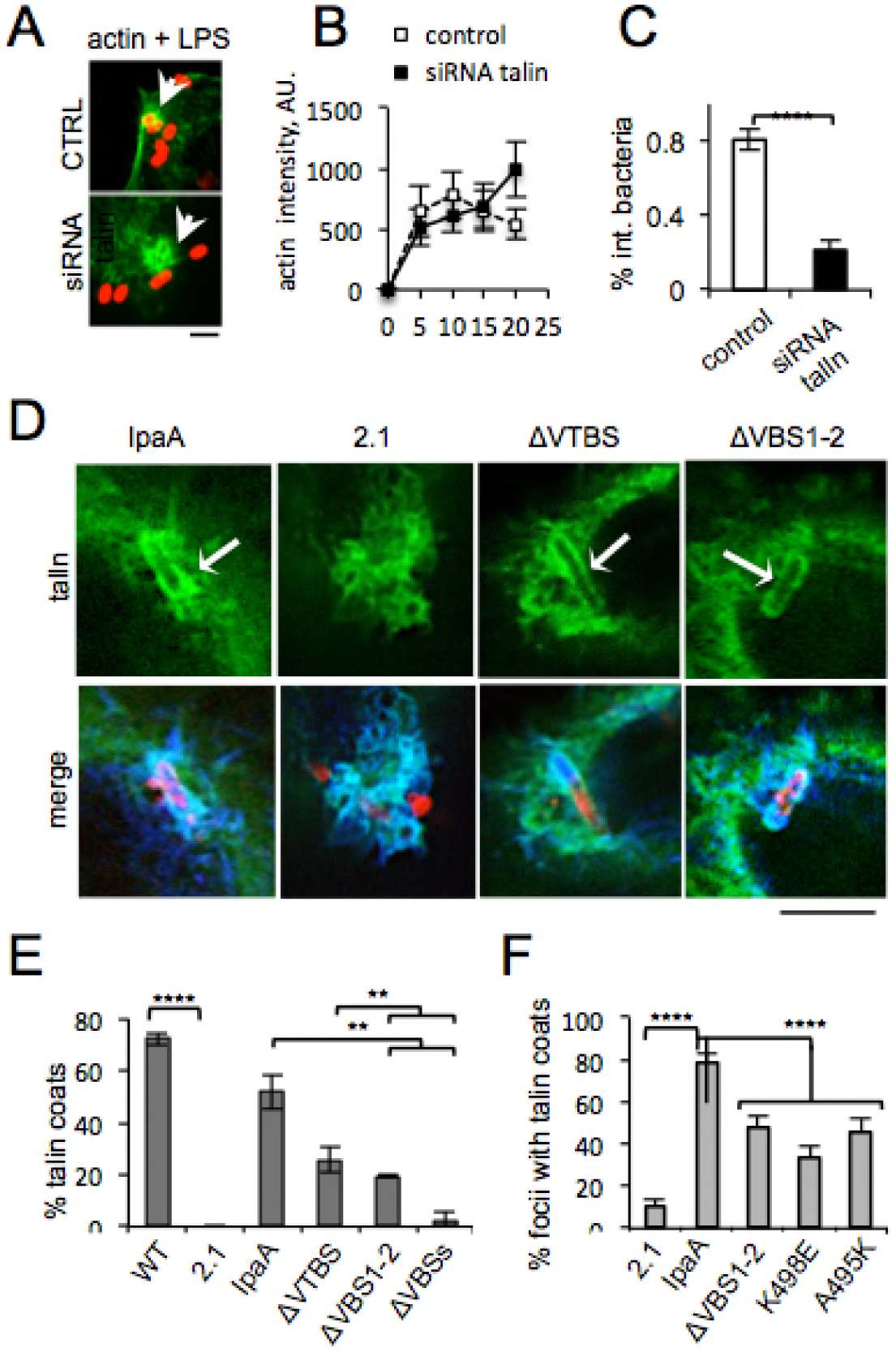
IpaA-dependent recruitment of talin is required for *Shigella* invasion. (A) Representative micrographs of control cells and cells treated with anti-talin siRNA prior to bacterial challenge at 37°C for 15 min. Green: actin; red: bacterial LPS. The arrows point to actin “cups” at bacterial cell contact (CTRL) or at the periphery of the foci (siRNA talin). Scale bar = 2 μm. (B) The average intensity of actin foci was determined at the indicated time points following bacterial challenge for control cells (empty squares) or anti-talin siRNA treated cells (black squares). N = 3, > 30 foci. (C) Cells were treated with anti-talin siRNA prior to bacterial challenge at 37°C for 15 min. The percentage of internalized bacteria after 30 minutes of infection with WT *Shigella* was determined (Methods). For each samples, n > 35 foci in at least three independent. Wilcoxon test. ****: p < 0.001. (D) Representative micrographs of cells were transfected with talin-GFP and challenged with *Shigella* strains for 30 min at 37°C. Red: bacterial; blue: actin; green: talin-GFP. Cells challenged with the *ipaA* mutant strain complemented with full length IpaA (Bricogne et al.), vector alone (2.1), IpaA ΔVTBS and IpaA ΔVBS1-2. The arrows point to talin coat structures surrounding invading bacteria. Scale bar = 5 μm. (E, F) Average percentage of actin foci forming talin-coat structures induced by the indicated bacterial strain ± SD (Methods). For each samples, n > 35 foci in at least three independent experiments. Chi Squared Test χ^2^ with post-hoc comparison (FDR correction for p-value).

These results indicate that talin is required for *Shigella* invasion and is recruited at entry sites in an IpaA-dependent manner to form actin coats surrounding invading bacteria. The direct talin binding activity of IpaA VTBS, however, appears dispensable for talin recruitment at invasion sites.

### Talin binding by IpaA VTBS is required for FA stabilization in infected cells

In addition to forming pseudo-adhesion structures in membrane ruffles during invasion, *Shigella* stabilizes FAs to prevent early detachment of infected cells (Sun et al., 2017). To investigate the role of IpaA VTBS in this process, cells were challenged with bacteria for 30 min and talin-containing adhesions were analyzed by immunofluorescence microscopy (Materials and Methods). As shown in Fig. 9, cells infected with the *ipaA* mutant complemented with full length IpaA remained spread and showed large adhesion structures (Figs. 9A-C). In contrast, cells challenged with *ipaA* mutant complemented with vector alone showed a significant decrease in size and number of adhesions (Figs. 9A-C). Complementation with IpaA ΔVBS1-2 containing VTBS restored adhesion structures although these latter were smaller than those observed for full-length IpaA (Figs. 9A, B). Strikingly, the A495K and K498E mutations affected the IpaA VTBS-dependent formation of adhesion structures at different degrees. Cells infected with *ipaA* / IpaA ΔVBS1-2-A495K were similar to cells infected with *ipaA* complemented with vector alone, with smaller and fewer adhesion structures compared to *ipaA* / IpaA ΔVBS1-2 (Figs. 9A-C). Complementation with IpaA ΔVBS1-2-K498E led to an intermediate phenotype with similar numbers of adhesions per cell but a clear shift in the distribution towards cells showing smaller adhesions (Figs. 9C, D).

**Figure 9.**
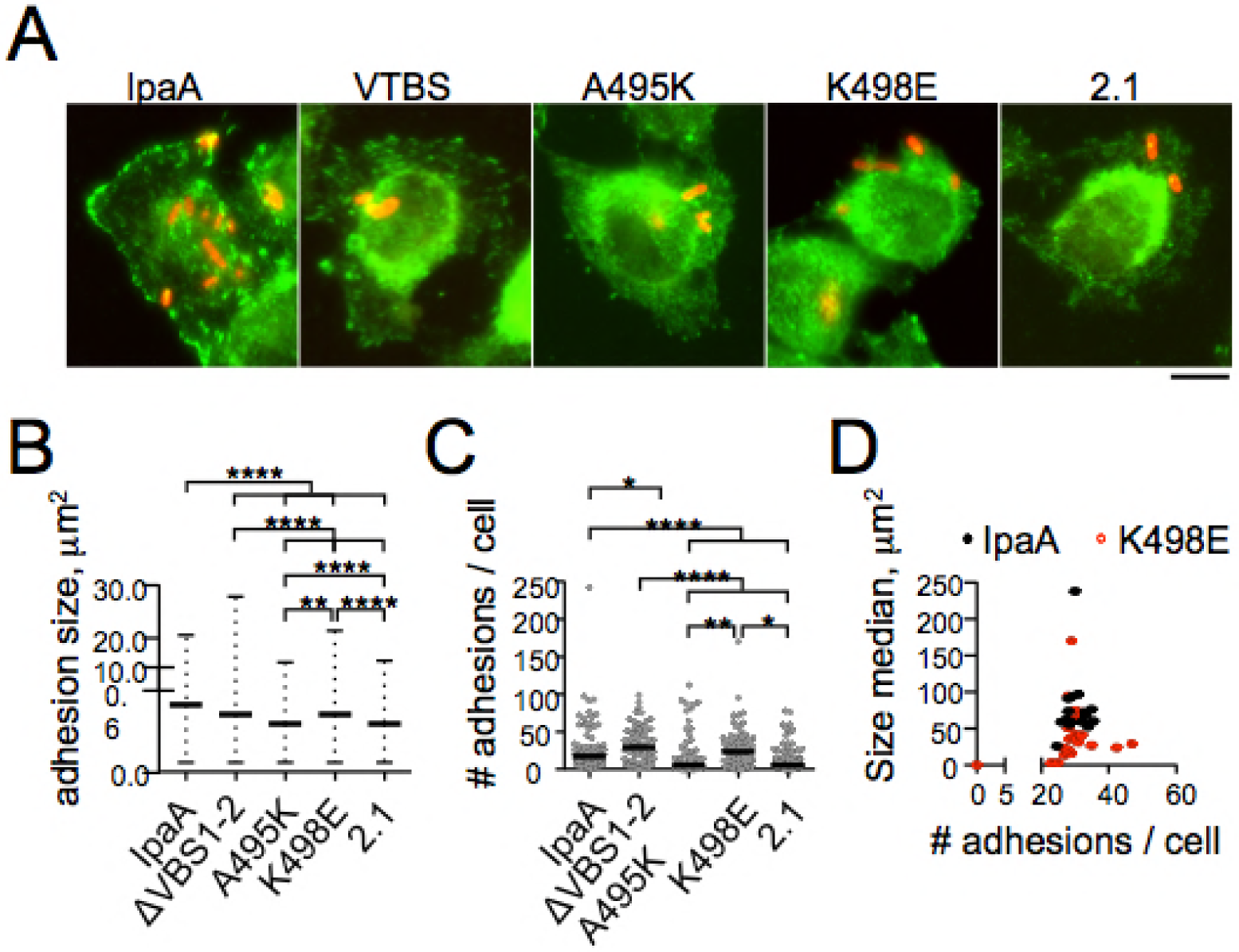
Talin-binding by IpaA VTBS is required for FA stabilization in *Shigella*-infected cells. HeLa cells were challenged with bacteria for 30 min at 37°C, fixed and processed forimmunofluorescence staining. Cells challenged with Shigella *ipaA* mutant complemented with: IpaA: full-length IpaA; ΔVBS1-2: IpaA Δ VBS1-2; A495K: IpaA Δ VBS1-2-A495K; K498E: IpaA Δ VBS1-2-K498E; 2.1: vector alone. (A) Representative micrographs. Red: bacteria; green: talin. Scale bar = 5 μm. (B, C) FAs were scored using automatic detection (Materials and Methods). The median adhesion size (B) and median number per cell (C) are indicated for each sample. (B) Median ± MAD: IpaA: 0.4949 ± 0.3535 (81 cells, N=3); ΔVBS1-2 0.4242 ± 0.2828 (60 cells, N=3); A495K 0.3535 ± 0.3535 (70 cells, N=3); K498E 0.4242 ± 0.2828 (75 cells, N=3); 2.1 0.3535 ± 0.2121 (111 cells N=3). (C) Median ± MAD: IpaA: 17 ± 11 (81 cells, N=3); ΔVBS1-2 29 ± 19 (60 cells, N=3); A495K 5 ± 5 (70 cells, N=3); K498E 23 ± 16 (75 cells, N=3); 2.1 5 ± 4 (111 cells N=3). (B) Kolmogorov-Smirnov test of probability distribution: **: p < 0.01; ****: p <0.001. (C) Kruskal-Wallis: *: p < 0.05; **: p < 0.01; ****: p < 0.001. (D) The median size of FAs is plotted for each cell as a function of the adhesions’ number per cell.

Together, these results are in line with the expression of ectopic IpaA VTBSs in transfection experiments and suggest that talin binding of IpaA VTBS plays an important role in stabilizing cell adhesions during *Shigella* infection.

## Discussion

Talin VBSs and RIAM or VLC1 / KANK TBSs are short amphipathic helices that target α-helical bundles in vinculin and talin (Gingras et al., 2005; Goult et al., 2013b; Zacharchenko et al., 2016). For these VBSs and TBSs, the interaction with the target bundle implicates a core of hydrophobic residues, often stabilized by electrostatic interactions between a pair of residues (Gingras et al., 2006; Goult et al., 2013b; Zacharchenko et al., 2016). Remarkably, these α-helices show high specificity for their respective targets, with TBSs targeting specific bundles in the talin rod and VBSs in general showing no binding to talin. A notable exception is RIAM TBS1, which, in addition to talin R2R3, was also shown to bind to vD1 in a manner similar to canonical activating VBSs (Goult et al., 2013b). The relevance of RIAM TBS1 binding to vinculin however, is not clear since its reported affinity for vinculin is about 10-fold lower than that of talin VBSs (Goult et al., 2013b). While talin bundles may look structurally similar, discrete changes Beyond the general rule of a hydrophobic core, the bundle specificity of TBSs / VBSs is determined by interactions that are unique for each interacting bundles and a-helices.

In this work, we report the first description of IpaA VTBS as a single α-helix that bind to vinculin as well as talin H1-H4 with high affinity, with a Kd of 54 pM and 174 nM, respectively. The apparent misbalance in affinity of IpaA VTBS in favor of vinculin is to be taken with caution, because ITC could not be used for its determination and surface plasmon resonance can lead to overestimated affinity. In addition, while IpaA VTBS requires talin stretching to bind to talin H1-H4, it has to compete out the vinculin tail interaction to induce its activation. Thus, IpaA VTBS may bind to talin or vinculin in an opportunistic manner, consistent with the observed preferential binding of GFP-IpaA VTBS to talin in filopodial adhesions, and to vinculin in mature adhesions.

Our sequence alignment provides a mechanistic basis for how IpaA VTBS acts as a dual talin-vinculin binder, a property that appears to be shared by talin H46. IpaA residues V499-L503-L507 are involved in interactions with talin or vinculin, suggesting that these constitute a common hydrophobic core essential for binding but not conferring specificity. Two other hydrophobic residues, IpaA I492-A495, however, appear to be specific for talin interaction. Also, electrostatic interactions involving residues IpaA R489 and K498 are important for talin specificity. Specifically, the lysine residue at position 498 in IpaA VTBS is critical for specific and high affinity binding to talin. This is explained by our structure showing IpaA K498 interacting with E621 in talin H1-H4 while not being involved in vinculin interaction.

We show that IpaA VTBS interacts with talin H1-H4 to form a compact bundle in sharp contrast to all other TBSs described to date that bind to unfolded talin H1-H4. In that perspective, IpaA VTBS mimics the talin H5 helix, both helices nesting identically in the hydrophobic groove and establishing similar polar interactions with talin H1-H4. IpaA VTBS, however, does not bind to talin H1-H5 indicating that as for other described TBSs, binding requires the removal of the H5 helix shown to occur upon activation and stretching of the talin molecule (Papagrigoriou et al., 2004). Stretching of talin requires its joint engagement with integrin receptors at the membrane and tethering to actin filaments subjected to retrograde flow or actomyosin contraction. Physical stretching of talin polypeptides using magnetic tweezers demonstrated the binding of vinculin upon unfurling of talin bundles and enabled to establish a hierarchy in the force-dependent exposure of VBSs in the various talin bundles (del Rio et al., 2009; Yao et al., 2016). For the talin R1, structural studies and molecular simulations indicate that vinculin binding requires unfurling of the bundle and full exposure of the VBS-H4 helix (Hytonen and Vogel, 2008; Papagrigoriou et al., 2004). Our structural data indicate that as opposed to vinculin, IpaA VTBS will not bind to-stretched talin H1-H4 expected in fully activated talin, but may target a talin conformer in which only the H5 helix will unfold from the H1-H4. There is no current experimental evidence for the existence of talin conformers containing partially stretched R1. Stretching on the talin R1-R3 domain using magnetic tweezers confirmed the unfolding at low forces of the R2 and R3 bundles, as expected from other studies, but failed to detect an intermediate unfolded state of the R1 bundle (Yao et al., 2014; Yao et al., 2016). The resolution limits of the technique, however, do not enable to detect displacement of a single helix H5, presumed to occur during IpaA VTBS binding to R1. Steered molecular dynamics on the other hand predicted that because of stronger hydrogen and salt bridges interactions between the talin H1 and H4 (talin VBS1) helices, applied force would result in a torque leading to dissociation the talin H5 helix from the H4 helix (Lee et al., 2007). SMD molecular dynamics also suggest a hierarchy in the unfolding of various talin VBS-containing bundles by force (Hytonen and Vogel, 2008). Specifically, talin VBS1 containing H1-H4 is expected to unfold at a low stretching force ranging from 15 to 25 pN (Yao et al., 2014). The labeling of semi-stretched talin by IpaA VTBS in filopodia, where forces of up to 15 pN are exerted (Romero et al., 2012), is fully consistent with this force range. The presence of a semi-stretched form of talin would explain the recruitment of talin and IpaA VTBS and the absence of vinculin at the filopodial distal extremity and in nascent adhesions, where talin is in a fully-stretched conformation.

Early studies had pointed to a role for talin in controlling filopodial elongation and retraction (Sydor et al., 1996). Since filopodial dynamics is controlled by actin polymerization at the tip of filopodia, these results are in line with a proposed regulatory role of talin on actin dynamics (Romero et al., 2012). In line with these studies, talin was since visualized as part of a MIT (Mig-10-RIAM-Lamellipodin (MRL)-Integrin-Talin) complex at the tip of filopodia in so-called “sticky fingers”, regulating their elongation (Lagarrigue et al., 2015). In the MIT complex, RIAM TBS1, through its binding to the talin R2 and R3 bundles, mediates the initial Rap1-dependent docking of inactive folded talin at the membrane and integrin activation (Chang et al., 2014; Goult et al., 2013b; Lee et al., 2013). Upon adhesion and stretching of talin, the R2R3 bundles are predicted to unfold first at low forces, promoting RIAM dissociation (Goult et al., 2013b; Yao et al., 2016). Because of the higher forces required for unfolding of the R1 bundle, IpaA VTBS likely targets a semi-stretched talin form following RIAM dissociation, although measurements of forces required to unfold the talin R2R3 bundles in the presence of RIAM are needed to confirm this.

We found VTBS expression to increase the length of filopodia in replating experiments. Because the stretching of talin by forces transduced by actin filaments requires its tethering to integrins adhering to the substrates, the talin conformer targeted by IpaA VTBS is not expected to directly promote the elongation of actin filaments as proposed for the MRL complex. Rather, IpaA VTBS may favor filopodial protrusion by stabilizing filopodial shaft adhesions. By forming a compact folded bundle with talin H1-H4, IpaA VTBS may stabilize a partially stretched talin conformer acting as a molecular clutch linking integrin receptors to filopodial actin filaments and adapted to the low force range exerted by these sensing organelles. Consistently, the IpaA A495K and K498E mutations that decrease talin-binding activity prevented the localization of IpaA VTBS in filopodial shaft adhesions and its effects on filopodia elongation. Because pathogens often mimic cellular processes, a similar role may be performed by talin H46 in the R10 bundle shown to unfold at intermediate stretching forces (Yao et al., 2014), which we show here to bind to talin H1-H4 with an affinity similar to IpaA VTBS. Unveiling of talin H46 in R10 may allow its binding to talin H1-H4 leading to talin-talin interaction. Talin intermolecular interactions may explain the clustering recruitment of talin in the absence of vinculin in distal filopodial shaft adhesions subjected to low pulling forces. As force increases, further stretching of talin will lead to the dissociation of talin VTBS-recipient bundle interaction and talin oligomers, to permit VBS-vinculin interactions, in line with the enrichment of vinculin at proximal filopodial shaft and basal adhesions.

Consistent with the reported relationship between filopodial and FAs (Hu et al., 2014; Partridge and Marcantonio, 2006), we found IpaA VTBS to label nascent adhesions. Interestingly, mutated IpaA VTBSs that are deficient for talin binding also failed to localize to FAs and did not promote their formation during bacterial infection. These results suggest that stabilization of the talin conformer by IpaA VTBS favors the formation of filopodial adhesions as well as FAs. Whether this talin conformer is involved in the maturation of filopodial adhesions into FAs is not clear because the latter may form in different modes (Hu et al., 2014; Partridge and Marcantonio, 2006; Rottner et al., 1999). FAs can be generated from the maturation and enlargement of basal adhesions supporting filopodial projections (Hu et al., 2014; Partridge and Marcantonio, 2006). Alternatively, FAs may result from the fragmentation and clustering of adhesions observed at the edge of migrating cells (Rottner et al., 1999). The mechanisms underlying these different modes of FA formation are not characterized, but may reflect the differential recruitment of cytoskeletal regulatory components in lamellipodia / lamellae and filopodial basal adhesions. Along these lines, we also observed different roles for IpaA VTBS at *Shigella* invasion sites and on cell adhesions. While talin is required for *Shigella* invasion and is recruited at entry sites in an IpaA-dependent manner, we found that its recruitment at entry foci does not require the talin-binding activity of IpaA VTBS. In contrast, the IpaA-mediated stabilization of FAs during bacterial infection requires talin-binding by IpaA VTBS, as indicated by the effects of cell challenge with bacteria expressing the IpaA VTBS A495K and K98E mutants. The reasons for this differential role of IpaA VTBS in bacterial-induced adhesion structures may relate to the difference observed for FA formation at the edge of migrating cells versus that during replating experiments. Indeed, *Shigella* invasion sites are characterized by actin polymerization and membrane ruffling, similar to what is observed at the cell migrating edge. It is possible that cytoskeletal proteins co-opted into membrane ruffles combined with vinculin activation by IpaA VBSs lead to talin recruitment. In contrast, adhesions forming at cell basal membranes during infection may critically depend on the recruitment of the talin conformer stabilized by IpaA VTBS.

## Material and Methods

### Antibodies

The anti-*Shigella* serotype V LPS rabbit antibody was described previously (Tran Van Nhieu et al., 1997)]. The mouse monoclonal anti-talin clone 8d4 antibody was from Sigma-Aldrich. Alexa 405 coupled anti-rabbit and Alexa 488 coupled anti-mouse IgGs were from Jackson Immunoresearch. Alexa 647-coupled phalloidin was from Invitrogen.

### Generation of expression constructs

Full-length human vinculin-mCherry (residues 1-1066) was generated by polymerase chain reaction (PCR) using 5’- CTGTCGACTGATGCCAGTGTTTCATACG-3’ / 5’- CTCCCGGGTCTGGTACCAGGGAGTCTTTC-3’ primers and cloned into the pmCherry-N1 (Clontech) vector using the *SalI* - *SmaI* sites. Full-length human talin fused to mCherry was from Addgene. Full-length human talin-GFP was a gift from K. Yamada. The A483 construct was PCR amplified using 5’- GGCGAATTCCCGGAGACACATATTTAACACG -3’ / 5’- GCCGTCGACTTAATCCTTATTGATATTCT -3’ primers and cloned into the *Eco*RI - *Sal*I sites of pGEX-4T-2 (GE Lifesciences). The GFP-A483 plasmid was generated by PCR amplification using the 5’-GCGATATCATGGCCAGCAAAGG-3’ and 5’- GCGCGGCCGCTTAATCCTTATTGATATTC-3’ primers and cloned into a pcDNA3.1 NT-GFP Topo (Invitrogen) following the manufacturer’s recommendations. The GFP-IpaAVTBS plasmid was generated by PCR amplification using the 5’-ACCCGGGGATTAAGCGGCC-3’ and 5’- ACCCGGGATCCTGATTTAGTTCC-3’ primers and the GFP-A483 plasmid as matrix. The amplicon was digested with *SmaI* and ligated using a T4 DNA Ligase. Mutagenesis to generate the GFP-IpaA VTBS A495K and K498E variants was performed using the following pairs of primers 5‘-GAGTTTGTTACTTTTTTTGATTTTTCAAATATCGTTTCCCGTG-3‘, and5‘-CGATATTTGAAGCTTCAAAAGAAGTAACAACAAACTCCCTAA-3’ and 5‘-TTAGGGAGTTTGTTGTTACTTCTTTTGAAGCTTCAAATATCG-3‘, respectively, using the Quickchange II site-directed mutagenesis procedure (Stratagene). All other enzymes used were from New England Biolabs. Plasmids containing talin VBS1 (H1-H4) residues 482-636 and GST-IpaA 524-633, as well as peptides IpaA-VBS1 (611-633) and VBS2 (565-586) were previously described (Izard et al., 2006; Papagrigoriou et al., 2004; Ramarao et al., 2007; Tran Van Nhieu and Izard, 2007). IpaA VTBS (N-TRETIFEASKKVTNSLSNLISLIGT-C, 488-512), and VTBS variant peptides K9498A (N-TRETIFEASKAVTNSLSNLISLIGT-C), K498E(N-TRETIFEASKEVTNSLSNLISLIGT-C), R489AK498A(N-TAETIFEASKAVTNSLSNLISLIGT-C) and A495K (N-TRETIFEKSKKVTNSLSNLISLIGT-C) were synthetized by Genscript USA Inc. Talin peptides VBS46 (N-YTKKELIECARRVSEKVSHVLAALQ-C, 1945-1970), Talin VBS6 (N-FQDVLMQLANAVASAAAALVLKAKS-C, 664-689), Talin VBS9 (N-RGVGAAATAVTQALNELLQHVKAH-C,765-789), Talin(N-NLKSQLAAAARAVTDSINQLITMCT-C,1330-1357) and Talin VBS50 (N-QVVLINAVKDVAKALGDLISATKAA-C, 2077-2102) were synthesized at the Scripps Institute proteomics facility (Jupiter Florida, USA). For crystallographic studies, the human talin H1-H4 domain (residues 481-636) preceded by an internal ribosome-binding site and a start codon was PCR-amplified and cloned into the *EcoRI* site of pET28a-IpaA-VBS3 (Park et al., 2011) to generate the bicistronic expression vector pET28a-IpaA-VBS3/VTBS-talinH1-H4. Plasmids were transformed into the *Escherichia coli* BL21(DE3) strain (Invitrogen). The binary complex of talin H1-H4·IpaA-VTBS was expressed in *E. coli* cells grown overnight in LB media containing kanamycin at a final concentration of 50 μg / ml at 30 °C.

### Yeast double hybrid analysis

The yeast two-hybrid analysis was performed using IpaA1-565 as bait to screen a human placental RP1 library, according to standard procedures and the Y2H protocole (Hybrigenics services).

### Protein purification

The talin H1-H4 was purified essentially as described (Borgon et al., 2004; Papagrigoriou et al., 2004). IpaA derivatives were purified by affinity chromatography in PBS (Phosphate Buffer Saline) using a GSTrap HP affinity column (GE Healthcare), cleaved with thrombin to remove the GST moiety, followed by size exclusion chromatography (HiLoad S200, Ge Healthcare). For crystallographic studies, talin H1-H4·IpaA-VBS3/VTBS binary complex was purified as described (Park et al., 2011). Protein concentration was determined using the BCA assay (Thermo Scientific). Samples were dialyzed in binding buffer and stored at −80°C at concentrations ranging from 1 to 10 mg/ ml.

### Native-PAGE analysis

Talin H1-H4 and IpaA peptides were incubated in binding buffer (25 mM Tris-HCl PH 7.0, 100 mM NaCl and 1 mM β-Mercaptoethanol) for 1hr at 4°C. After incubation, samples were resuspended in 2 x Native loading buffer (62,5mM Tris pH 6,8 containing 25% Glycerol) and separated by Tris-Glycine Native-PAGE electrophoresis (Schagger et al., 1994). Gels were stained using standard colloidal Coomassie stain.

### SEC-MALS

The purified proteins IpaA483-633, IpaA524-633 and talin H1-H4 were used at 20μM equimolar concentrations and incubated at 4°C for one hour in binding buffer (25 mM Tris-HCl PH 7.0, 100 mM NaCl and 1 mM β-Mercaptoethanol). 200μls of the protein mixtures were analyzed by size-exclusion chromatography (SEC) on a Superdex 200 10/300 GL (GE Healthcare) using a Shimadzu Prominence HPLC. Multi-angle laser light scattering (MALS) was measured with a MiniDAWN TREOS equipped with a quasi-elastic light scattering module and a refractometer Optilab T-rEX (Wyatt Technology). Protein concentration was determined using a specific refractive index (dn/dc) of 0.183 at 658 nm.

### Crystallization, structure determination, and crystallographic refinement

Crystals of the IpaA-VTBS / talin H1-H4 complex were obtained by mixing 1 µl of protein complex at 7 mg/ml in buffer 25 mM Tris pH 7.4 and 150 mM NaCl with 1 µl of 0.2 mM ammonium sulfate, 0.1 mM sodium acetate pH 4.6 and 30 % PEG 4000 by vapor diffusion at 292 K. Crystals were cryoprotected with a solution consisting of the reservoir supplemented with 20 % v/v ethylene glycol and flash frozen. X-ray diffraction data were collected at 100 K on a single crystal at PROXIMA-1 beamline at the SOLEIL synchrotron (Saint-Aubin, France). Data were indexed and processed using XDS^25^ and corrected for anisotropy with the STARANISO server (staraniso.globalphasing.org). Structure solution was obtained by molecular replacement with Phaser^26^ using as search template a monomer of talin-H1-H4 derived from the pdb entry 1sj8 lacking residues 515 to 538. A clear solution was obtained (TFZ = 28.5; LLG = 2105.5) Refinement was done with autoBUSTER^27^ using NCS and alternating with manual building in Coot^28^. One TLS group per chain was used at the end of the refinement.

### Isothermal titration calorimetry

Protein interaction was analyzed by microcalorimetry using an ITC200 calorimeter (MicroCal) at 25°C. 200 μls of 20-100 μM of talin H1-H4 protein in binding buffer (25 mM Tris-HCl pH 7.0, 100 mM NaCl and 1 mM β-mercaptoethanol) were added to the cell and binding was measured in the presence of different concentrations of IpaA / talin VBSs peptides. In other experiments, A483 or A524 were added to the cell and binding of talin H1-H4 (100 μM) was measured. Typically 20-40 injections of 2 μls of ligand were made with intervals of 320 seconds between each addition, with a reference power of 12 μcal / sec. Data was analyzed using the MicroCal software provided by manufacturer.

### Cell culture and bacterial strains

HeLa cells (ATCC) were incubated in RPMI (Roswell Park Memorial Institute) medium containing 5% FCS (fetal calf serum, Gibco^®^) in an incubator with 5% CO_2_. Mouse Embryo Fibroblast cells (MEF, ATCC) were incubated in DMEM (11965092, GIBCO) medium containing 10% FCS (fetal calf serum, Gibco^®^) in an incubator with 5% CO_2_. The wild type *Shigella flexneri*, isogenic mutants, and complemented *ipaA* mutant strains, as well as wild type *Shigella* expressing the AfaE adhesin were previously described (Park et al., 2011). Bacterial strains were cultured in trypticase soy broth (TCS) medium at 37°C. When specified, antibiotics were added at the following concentrations: carbenicillin 100 μg/ml, kanamycin 20 μg/ml.

### siRNA transfection

HeLa cells were seeded at a density of 10^5^ cells in wells containing a 22 ×; 22 mm coverslip in a 6-well plate. The following day, cells were transfected with of anti-talin 1 siRNA (Stealth Select RNAi, catalogue no. 1299003, Invitrogen, oligo 804; sequence: 5’-CCAAGAACGGAAACCUGCCAGAGUU-3’) or anti-vinculin siRNA (Stealth Select RNAi, catalogue no. 1299001, Invitrogen, oligo VCLH55111259) duplex at the indicated concentrations and time periods.

### Cell challenge with *Shigella* strains

HeLa cells seeded at 2 ×; 10^5^ cells in coverslip-containing 33 mm-diameter wells the day before, were transfected with the indicated constructs using the JetPei^®^ transfection reagent. After 16 hours, cells were challenged with *Shigella* strains coated with poly-L-lysine, as previously decribed (Park et al., 2011).

### Immunofluorescence microscopy analysis

For GFP-IpaA VTBS and talin / vinculin co-localization experiments MEF cells were seeded at a density of 1 ×; 10^4^ cells in a µ-Dish 35 mm 15 kPa stiffness chamber (Cat #81391, ibidi) coated with 10 µg / ml of Fibronectin (Calbiochem). Cells were transfected with plasmids encoding human talin-mCherry, vinculin-mCherry and GFP-IpaA VTBS using the Lipofectamine 2000 transfection reagent (Life Technologies). For replating experiments HeLa cells co-transfected with vinculin-mCherry and talin-GFP, were resuspended by trypsinization and replated on glass coverslips. After 20 minutes the samples were fixed, processed for immunofluorescence microscopy, and mounted on slides using Dako mounting medium (Dako, Agilent Technologies), as described (Romero et al., 2012).

Samples were analyzed using an Eclipse Ti microscope (Nikon) equipped with a 100x objective, a CSU-X1 spinning disk confocal head (Yokogawa), and a Coolsnap HQ2 camera (Roper Scientific Instruments), controlled by the Metamorph 7.7 software. The percent of internalized bacteria was scored as one if the number of internalized bacteria per foci was at least the 50% of the total number of bacteria in the foci or zero if below, in three independent experiments (n_control_=27, n_siRNATln_=23). For control and anti-talin siRNA transfected cells, the percent of internalized bacteria was compared using a Chi-Squared test (R Statistical Software).

### Image processing and analysis

Analysis was performed in Icy Bioimaging Analysis software (de Chaumont et al., 2012). Scans of GFP-IpaA VTBS, Vinculin-mCherry and Talin-mCherry were performed as follows. Defined ROIs in median projections of basal in-focus planes were segmented in concentric areas, taking into account the edge of the cell. Average values of intensities were normalized to the maximum and minimum average intensities of the ROI. For talin-GFP and vinculin-mCherry filopodial distribution, Saturated sum projections of images from 20 minutes-replated cells were used for Concentric Area Scan analysis. For the quantification of the number of FAs, a semi-automated protocol was developed using Icy software (de Chaumont et al., 2012). Spinning-disk fluorescent microscopy planes were used to detect GFP-IpaA VTBS structures using HK means thresholding and overlaid binary masks obtained from the threshold projections of F-actin labeled images (Max-entropy method). FAs were detected as spots positive for both GFP-IpaA VTBS and actin structures using Wavelet Spot Detector. FAs were detected as spots positive for both GFP-IpaA VTBS above background intensity levels. Filopodia length and number were determined manually from actin-labeled projections. IpaA VTBS, talin, vinculin and vD1 filopodial clusters were identified using Wavelet Spot Detector in Saturated sum projections. Co-localizing clusters were determined by overlaying the separate detections. Distance to cell body was determined using ROI Inclusion analysis plugin.

### Statistical Analysis

Bacterial internalization and talin coat-structure formation were analyzed a contingency table using in a Pearson Chi-Squared test (R Statistical Software). A Post-hoc pairwise Chi-square test (NCStats package, R Statistical Software) with FDR p-value correction was further used to compare the distribution between the various strains. n > 35 foci, N = 3. The correlation of the intensity of talin and vinculin labeling was analyzed using a Wilcoxon rank sum test (R Statistical software).

## Acknowledgments

The authors thank Dr. Olivera Francetic for careful reading of the manuscript and Dr. David Stroebel (IBENS) for his help with ITC measurements. This work benefited from the facilities of the *Plateforme de Mesures d’Interaction des Macromolécules* (PIM), I2BC (http://www.cgm.cnrs-gif.fr/) and from the *Plateforme de Proteomique (Ecole Normale Superieure)*. This work was supported by grants from the Inserm, the CNRS and the Collège de France to the CIRB, as well as grant from the PSL Idex project “Shigaforce”. CV-G is a recipient of a PhD fellowship from the Memolife Labex and a post-doctoral fellowship from the PSL Idex. NC is a recipient of an IRG Marie-Curie fellowship. DA is funded by CONACYT and the Labex Memolife international PhD program.

## Author contributions

DA, NC, NQ-D, LP, HJ P performed and analyzed experiments. MF, and TI analyzed experiments. C V-G, C B-N, and GTVN performed, analyzed experiments and wrote the manuscript.

## Expanded View Information

**Expanded View Table 1.** Results of the Yeast two-hybrid screen of a human placental RP1 cDNA library using IpaA1-565 as bait (Hybrigenics SA, France). Certitude: Level of confidence: A, very high; E, prey domains connected 6 times.

| Clone No. | Gene | Start...Stop | Frequency | Certitude |
| --- | --- | --- | --- | --- |
| 1 | hTLN1 | 1446..2324 | 2 | A |
| 2 | hTLN1 | 1446..2329 | 2 | A |
| 3 | hTLN1 | 5118 | 1 | A |
| 4 | hTLN1 | 5268..6788 | 1 | A |
| 5 | hTLN1 | 5358 | 8 | A |
| 6 | hTLN1 | 5358..6787 | 14 | A |
| 7 | hTLN1 | 5379..6782 | 2 | A |
| 8 | hTLN1 | 5457 | 1 | A |
| 9 | hTLN1 | 5457..6790 | 1 | A |
| 10 | hTLN1 | 5558..6913 | 1 | A |
| 11 | hTLN1 | 5865..6785 | 1 | A |
| 12 | hTLN1 | 5871 | 1 | A |
| 13 | hTLN1 | 5871..6787 | 24 | A |
| 14 | hTLN1 | 5871..6789 | 1 | A |
| 15 | hTLN1 | 5871..6790 | 3 | A |
| 16 | hTLN1 | 5871..6800 | 1 | A |
| 17 | hTLN1 | 5872..6787 | 1 | A |
| 18 | hTLN1 | 5895..6788 | 1 | A |
| 19 | hTLN1 | 5919 | 1 | A |
| 20 | hTLN1 | 5919..6785 | 1 | A |
| 21 | hTLN1 | 5928..6785 | 1 | A |
| 22 | hTLN1 | 5982..6788 | 1 | A |
| 23 | hTLN1 | 6297..6814 | 2 | A |
| 24 | hTLN1 | 6414..6786 | 5 | A |
| 25 | hTLN1 | ..6781 | 1 | A |
| 26 | hTLN1 | ..6784 | 3 | A |
| 27 | hTLN1 | ..6785 | 3 | A |
| 28 | hTLN1 | ..6786 | 7 | A |
| 29 | hTLN1 | ..6789 | 1 | A |
| 1 | hVCL | 20..1505 | 5 | E |
| 2 | hVCL | 96..987 | 2 | E |
| 3 | hVCL | ..986 | 2 | E |
| 4 | hVCL | ..1115 | 1 | E |

**Expanded View Table 2.** Summary of data collection and refinement statistics.

| Talin H1-H4 / IpaA VTBS |  |
| --- | --- |
| **Data collection** |  |
| Space group | P 21 21 21 |
| Wavelength (Å) | 0.9786 |
| Cell dimensions |  |
| *a*, *b*, *c* (Å) | 66.26, 96.36, 175.69 |
| *a*, *b*, *g* (°) | 90, 90, 90 |
| Resolution (Å) | 48.18 - 2.50 (2.59-2.50) |
| *R*<sub>merge</sub> | 0.083 |
| *R*<sub>meas</sub> | 0.094 |
| *I* / *σ**I* | 9.4 (0.7) |
| *CC*<sub>1/2</sub> | 0.998 (0.567) |
| Completeness (%) | 99.2 (95.7) |
| Redundancy | 4.8 (4.8) |
| **Refinement** |  |
| Resolution (Å) | 36.33 – 2.50 |
| No. reflections | 39441 |
| *R*<sub>work</sub> / *R*<sub>free</sub> | 0.196 / 0.228 |
| No. atoms |  |
| Protein | 7167 |
| Water | 140 |
| *B*-factors |  |
| Protein |  |
| Talin H1-H4 | 69.06 |
| IpaA VTBS | 67.52 |
| solvent | 54.0 |
| R.m.s. deviations |  |
| Bond lengths (Å) | 0.010 |
| Bond angles (°) | 1.09 |
| *Data were collected on a single crystal. Values in parentheses are for highest-resolution shell. |  |

**Expanded View Figure 1.**
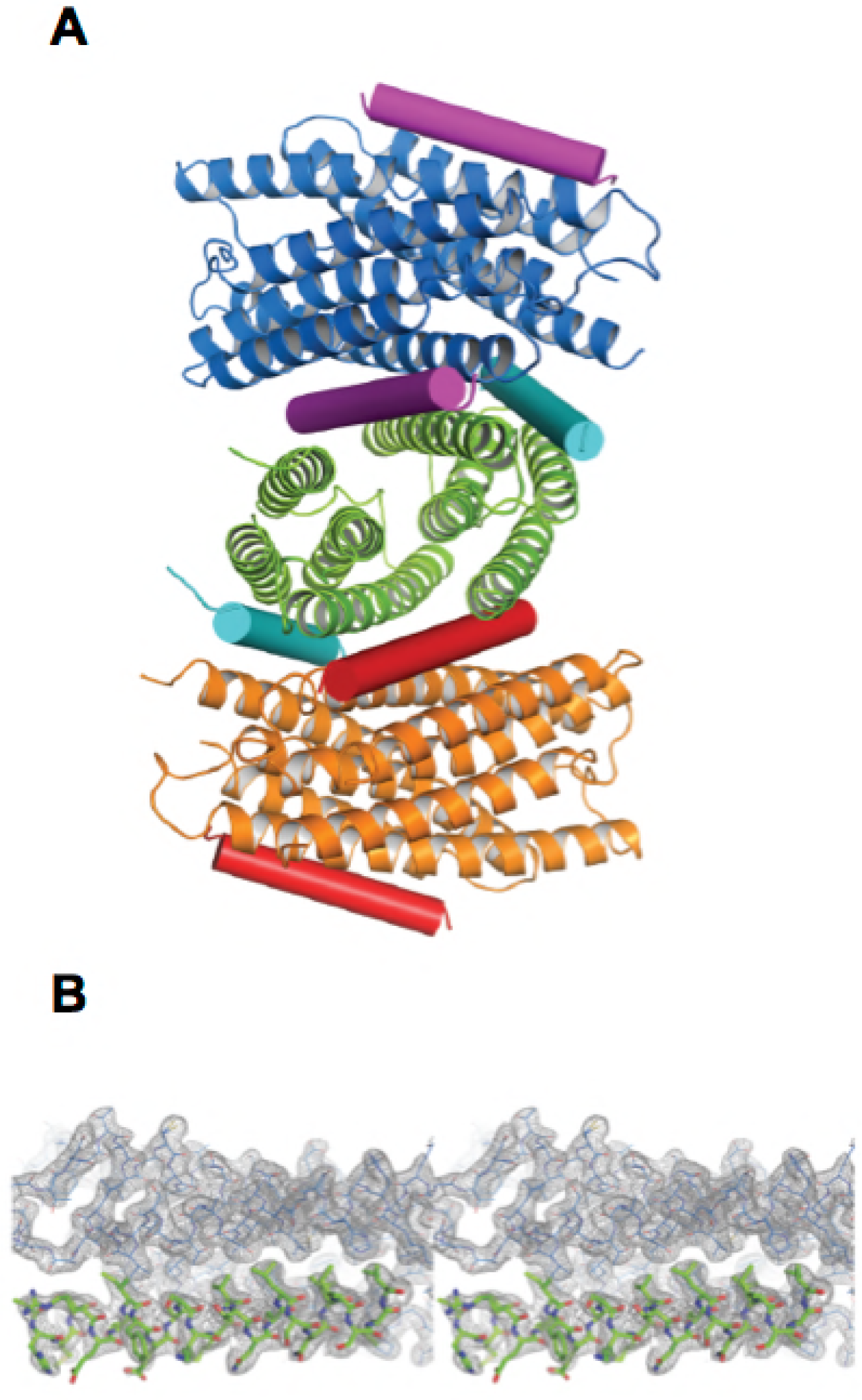
(A) Crystallographic unit cell of talin H1-H4 bound to IpaA VTBS. The three dimers of talin H1-H4 are represented in cartoons colored in blue, green and orange and the bound IpaA VTBS peptides are shown as cylinder in purple, cyan and red. **(B)** Stereoview of the 2mF_0_-DF_c_ map contoured at 1σ showing the electron density of one IpaA VTBS peptide, in green, interacting with H2 (top view) and H4 (back view).

**Expanded View Figure 2.**
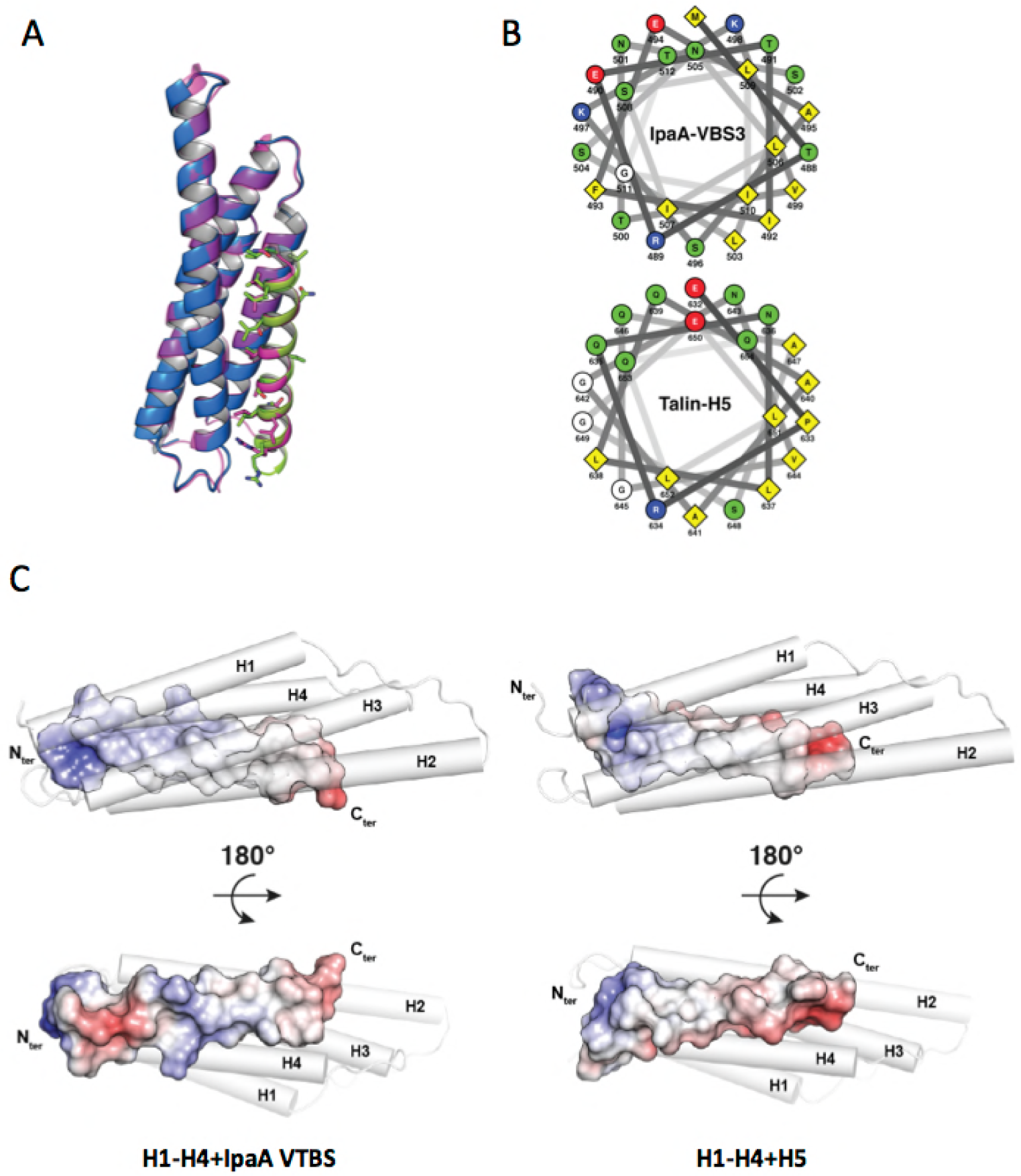
IpaA VTBS mimics talin H5. (A) Superimposition of talin H1-H4:IpaA VTBS and talin H1-H5 structures. Talin H1-H4 bound to IpaA VTBS (green) and to talin H5 (pink) are shown as a cartoon in blue and magenta, respectively. (B) Helical wheel analysis of IpaA VTBS (top) and talin H5 (bottom) indicates that both are amphipathic α-helices with conserved residues in their hydrophobic interphase. The hydrophobic residues, polar residues, positively and negatively charged residues are colored yellow, green, blue, and red, respectively. (C) Comparison of talin H1-H4 binding to IpaA VTBS and talin H5. Talin H1-H4 is shown in white cylinder while the electrostatic surface of IpaA VTBS and H5 are represented as calculated with APBS.

**Expanded View Figure 3.**
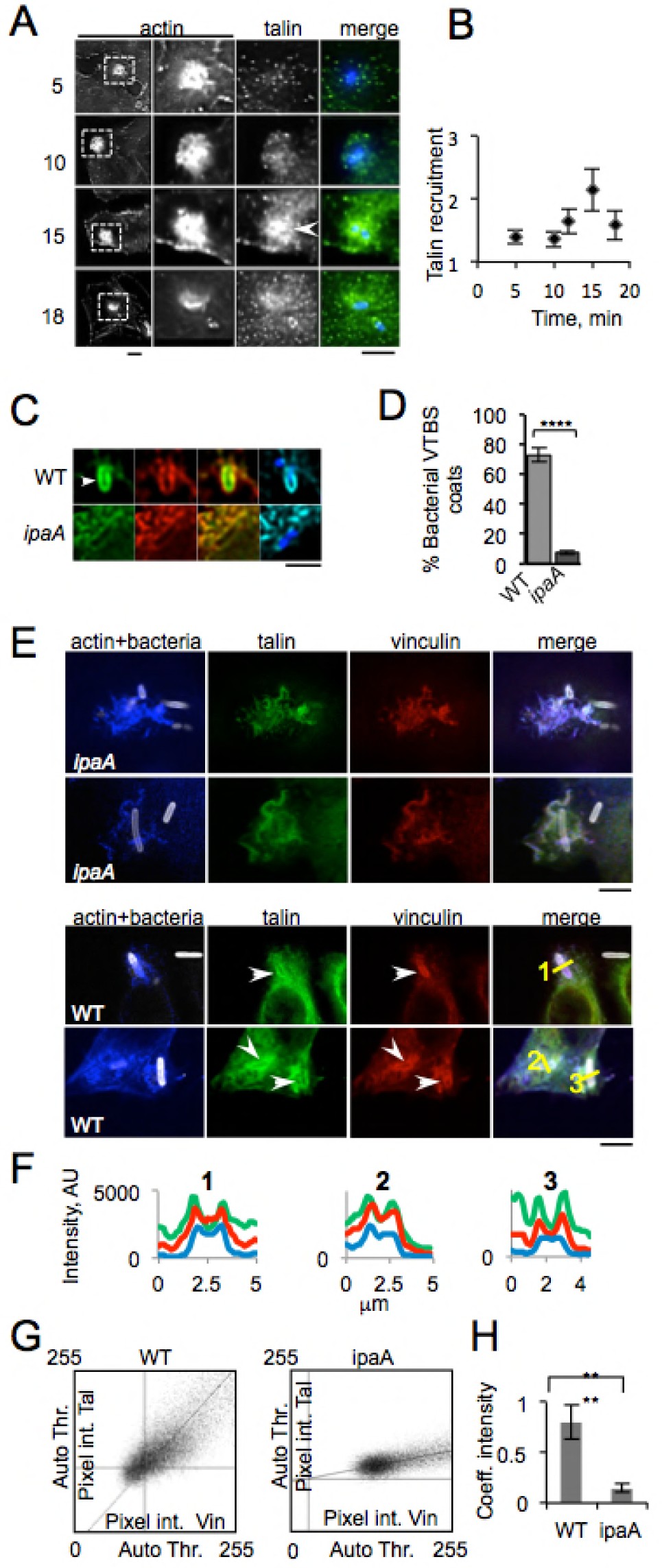
Talin and vinculin co-localize at IpaA-dependent coat structures. **(A, B)** HeLa cells were challenged with wild-type *Shigella*. Samples were fixed and processed for immunofluorescent staining of bacterial LPS (blue), talin (green). (A) Representative micrographs of cells challenged with WT *Shigella* for the time indicated in min on the left, with a larger magnification of the inset boxed in the actin staining in the left panel. The arrows point to talin coalescing around invading bacteria. Scale bar = 5 μm. (B) Average intensity of talin staining at actin foci was quantified and the recruitment expressed as a ratio relative to a corresponding cell control area (Methods). For each samples, n > 30 foci in at least three independent experiments (C, D) Cells co-transfected with GFP-IpaA-VTBS and talin-mCherry were challenged with WT *Shigella* or the *ipaA* mutant. (C) Representative micrographs. Green IpaA VTBS; red: talin; blue: bacteria; cyan: F-actin. Scale bar = 5 μm. Talin and GFP-IpaA VTBS are detected at the filopodial tip where bacterial capture occurs, as well as in bacterial coats in invasion foci induced by WT *Shigella* but not the *ipaA* mutant. (D) Percentage of invasion foci showing GFP-IpaA VTBS labeled bacterial coats. N = 2, > 60 foci. Wilcoxon test. ****: p < 0.001. (E-H) Talin-GFP and vinculin-mCherry transfected cells were challenged with *Shigella* strains for 30 min at 37°C. Samples were processed for immunofluorescence staining. For each sample, more than thirty stack fields were acquired for each condition in two independent experiments. The results are expressed as the fluorescence intensity ratio of Talin between the foci and the cytoplasm (R) using the following formula: R = I_F_-I_o_/ I_c_-I_o_, with **I_F_**: Medium intensity of talin at the entry foci; **I_C_**: Average intensity of talin in the cytoplasm of the infected cell; **I_O_**: Medium intensity of background in a region containing neither cells nor bacteria. To correlate of the intensity of talin and. vinculin labeling, regions were drawn around bacteria and slope of the correlation of the pixel intensity was determined using the Correlation Threshold plug-in (ImageJ) (WT: n = 33 foci, N = 3; *ipaA* mutant: 18 foci, N = 3) (A, B) actin: red; talin: green; vinculin: red. (E) Representative micrographs of samples challenged with the indicated bacterial strains (grey). Z-project: reconstruction of merge labeling. Right panels: single confocal planes corresponding to the magnified inset in the left panels. Scale bar = 5 µm. (F) Normalized fluorescence intensity scans of the lines depicted in panel A. (G) Scatter plots showing the intensity correlation of pixels with vinculin and talin co-localization at bacterial coats for WT and *ipaA* mutant strains. (H) The median of the regression coefficient of the intensity of talin and vinculin around the bacteria at WT and *ipaA* foci (n_WT_ =33 and n_ipaA_ =18) were measured as 0.80 ± 0.16 and 0.14 ± 0.04 respectively, and compared using a Wilcoxon rank sum test. ****: p < 5 x10^−9^.

